# Learning, sleep replay and consolidation of contextual fear memories: A neural network model

**DOI:** 10.1101/2025.06.20.660661

**Authors:** Lars Werne, Angus Chadwick, Peggy Seriès

**Affiliations:** Center for Doctoral Training: Biomedical AI, University of Edinburgh, Edinburgh, United Kingdom; Institute for Adaptive and Neural Computation, University of Edinburgh, Edinburgh, United Kingdom

## Abstract

Contextual fear conditioning is an experimental framework widely used to investigate how aversive experiences affect the valence an animal associates with an environment. While the initial formation of associative context-fear memories is well studied – dependent on plasticity in hippocampus and amygdala – the neural mechanisms underlying their subsequent consolidation remain less understood. Recent evidence suggests that the recall of contextual fear memories shifts from hippocampal-amygdalar to amygdalo-cortical networks as they age. This transition is thought to rely on sleep. In particular, neural replay during hippocampal sharp-wave ripple events seems crucial, though open questions regarding the involved neural interactions remain.

Here, we propose a biologically informed neural network model of context-fear learning. It expands the scope of previous models through the addition of a sleep phase. Hippocampal representations of context, formed during wakefulness, are replayed in conjunction with cortical and amygdalar activity patterns to establish long-term encodings of learned fear associations. Additionally, valence-coding synapses within the amygdala undergo overnight adjustments consistent with the synaptic homeostasis hypothesis of sleep. The model reproduces experimentally observed phenomena, including context-dependent fear renewal and time-dependent increases in fear generalisation.

Few neural network models have addressed fear memory consolidation and to our knowledge, ours is the first to incorporate a neural mechanism enabling it. Our framework yields testable predictions about how disruptions in synaptic homeostasis may lead to pathological fear sensitization and generalisation, thus potentially bridging computational models of fear learning and mechanisms underlying anxiety symptoms in disorders such as PTSD.

**Author Summary:** How do we learn to fear certain environments? Why do some fear memories fade while others persist or even grow stronger over time? Scientists have long used laboratory experiments to study how animals associate danger with a particular context. These studies have helped identify brain regions involved in fear learning, including the amygdala, hippocampus, and cortex, and have inspired many computational models of how fear is acquired in the brain.

However, most models focus only on what happens when fear is first learned, overlooking how these memories evolve in the days that follow and the role of sleep in this process. In this work, we present a neural network model that captures how fear memories are strengthened or reshaped during sleep. It builds on earlier models by incorporating memory replay and synaptic homeostasis, two brain processes believed to support emotional memory consolidation. Our model identifies neural processes that help make fear memories persistent, suggests that sleep is necessary to maintain adaptive behaviour after threatening experiences, and proposes that sleep disruptions mediate the harmful impact of stress on emotional regulation. By extending amygdala-based models of fear learning to include post-learning dynamics, our work offers new insight into how emotional memories are stabilised.

## Introduction

Despite robust biological evidence of sleep’s essential role in emotional memory consolidation, computational models of fear learning typically neglect sleep-dependent processes. We propose a neural network model that incorporates sleep-mediated replay and synaptic homeostasis, offering a new mechanistic perspective on fear memory formation, retention, and generalisation.

Pavlovian Fear Conditioning, in which an initially neutral cue or context is repeatedly paired with an aversive stimulus (US), is a fundamental paradigm for investigating the neural mechanisms underlying fear learning [1]. In *contextual* fear conditioning, fear becomes associated with the combined environmental and internal elements defining the conditioned *context* [2]. Understanding how contextual fear memories form, persist, and generalize is crucial both for basic neuroscience and for clinical conditions characterized by maladaptive fear, such as Post-Traumatic Stress Disorder (PTSD) [3, 4].

Neural mechanisms underlying fear memory formation extend beyond awake experiences into subsequent periods of sleep. During sleep, neural replay occurs simultaneously across hippocampal and neocortical circuits, reflecting activity patterns learned during prior experiences [5, 6]. This coordinated replay has robust experimental support and is critically involved in the consolidation of associative memories, including those formed through fear conditioning [7]. The basolateral amygdala (BLA), long recognized for its central role in emotional learning [8], also exhibits activity coordinated with hippocampal replay during sleep. Although open questions regarding precise mechanisms and functional significance remain, emerging evidence suggests that coordinated replay across hippocampus and amygdala may help bind contextual representations to emotional salience, driving emotional memory retention [9].

Despite these findings, existing computational models of associative fear learning [10–12], originating from classical frameworks such as the Rescorla-Wagner model [13–15], typically omit these sleep-dependent consolidation mechanisms. These models assume immediate and stable updating of emotional associations during awake learning episodes. As a result, they struggle to account for clinical observations where maladaptive fear symptoms — e.g., in PTSD — often emerge *gradually*, rather than immediately, after intense fearful, or traumatic, experiences. This discrepancy implies a fundamental limitation: by omitting critical neural events occurring after initial learning episodes, existing models may inherently be unable to explain the gradual onset of pathological fear.

To overcome this gap, we propose a new neural network model of contextual fear learning that explicitly incorporates sleep-dependent replay and synaptic homeostasis. Our simulations demonstrate how disruptions in these sleep-related processes – e.g., under psychological stress – could lead to amygdala hyperactivity, enhanced fear acquisition and heightened generalisation. While formulated at a fairly high level of abstraction, our work not only provides a qualitative framework linking sleep to emotional memory processing but also lays a foundation for further computational studies and experimental investigations into fear learning and its clinical implications.

## Methods: Model description

To investigate interactions between sleep processes and fear learning, we developed a computational model that combines a hippocampal sleep replay mechanism, proposed by Fiebig et al. for systems consolidation of episodic memories [16], with an amygdalo-centric framework for associative fear learning, inspired by previous neural network models [11, 17]. By drawing from these computational approaches, we provide a novel, mechanistic account of how context fear memories transition from transient hippocampal-amygdalar representations into stable amygdalo-cortical associations [18].

Our model reproduces key behavioural phenomena observed in fear conditioning – including context-dependent fear renewal [19] and enhanced fear learning under stress [20, 21]. Its architecture is sketched in Fig 1. Environmental inputs (or *contexts*) are represented as binary activity patterns within the sensory cortex (SC) module, from where they are transmitted to each of our model’s three primary regions [22–24]:

**Fig 1.**
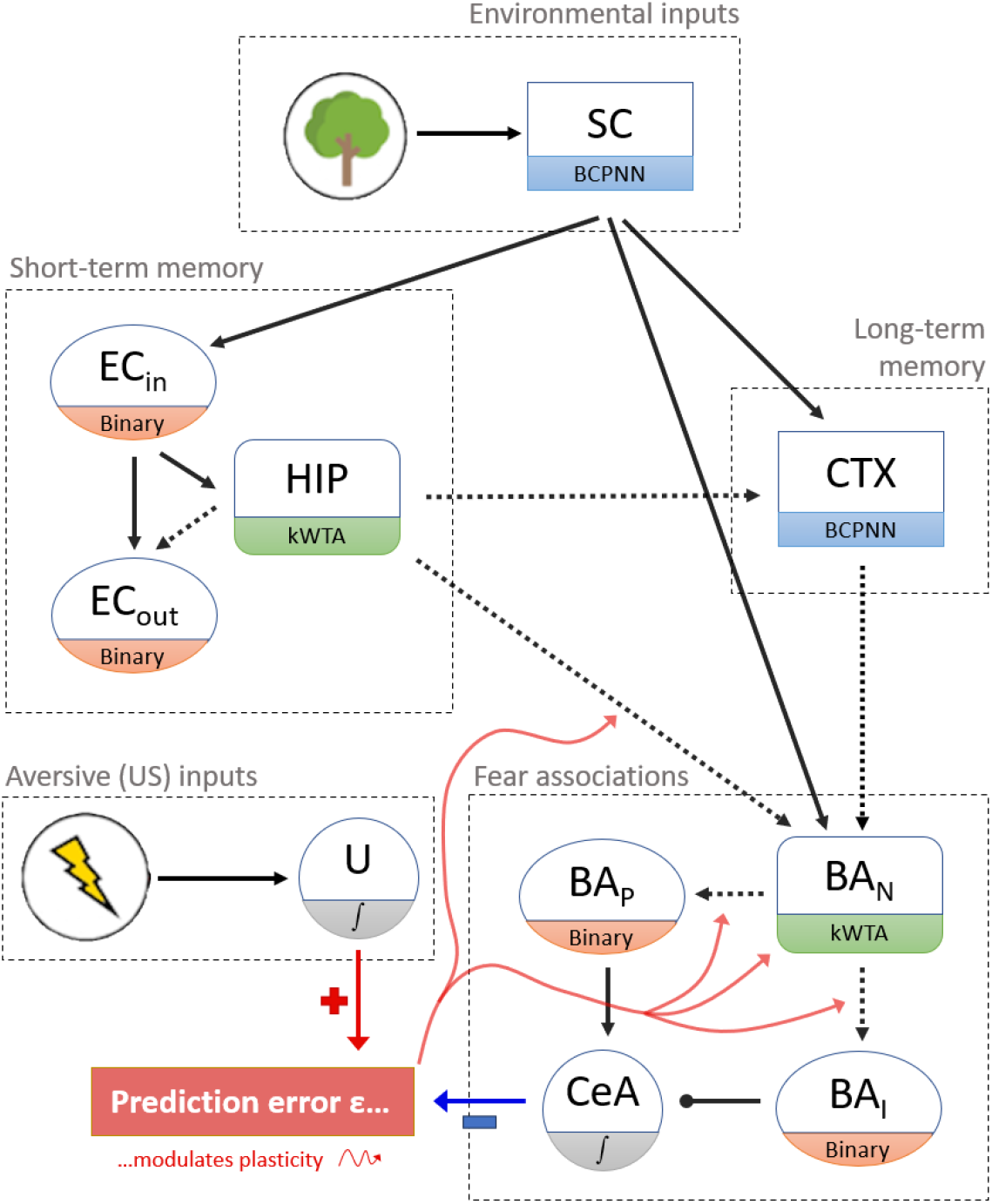
Model sketch. Nodes of the graph correspond to *modules*. Dotted/solid arrows denote the existence of plastic/non-plastic synapses between modules. During *Perception*, environmental inputs are passed to the *engram* modules HIP, CTX and BA_N_, activating patterns that may be remembered via Hebbian plasticity, so that the model’s state may converge onto them during later *Recall*. To allow either fear or safety to be associated with a context, synapses from HIP onto BA_N_ are strengthened when a prediction error occurs. Prediction errors are defined as the difference between the current US input and fear response (CeA output). Errors promote Hebbian plasticity within BA_N_ and on HIP *→* BA_N_ synapses, allowing the context to be associated with valence. Positive / negative errors further strengthen synapses from BA_N_ onto BA_P_ / BA_I_, increasing / decreasing the amount of fear associated with the current environment. Nodes labelled with an S-shape are single units. All other modules are sets of neurons – the BCPNN (Bayesian Confidence Propagation Neural Network), kWTA (k-Winner-Takes-All) and ‘binary’ module types are defined in S1 Appendix. **Abbreviations:** SC = Sensory Cortex. EC_(in / out)_ = Entorhinal Cortex (Input- / Output layers). HIP = Hippocampus. CTX = Neocortex. BA_(**N** / **P** / **I**)_ = Basal Amygdala (valence-**N**eutral / fear-**P**romoting / fear-**I**nhibiting sets of neurons). CeA = Central Nucleus of Amygdala.

- **Hippocampus (HIP)**, which rapidly encodes and temporarily stores context information using Hebbian learning. During sleep, recurrent excitation and fast inhibitory processes drive the replay of these transient engrams.
- **Neocortex (CTX)**, which forms long-term representations through slow synaptic updates. Sleep replay reinforces and stabilizes cortical traces, allowing them to maintain fear associations independently of hippocampal support.
- **Amygdala (BA & CeA)**, which associates contexts with valence. Context identity, fear, and safety are encoded by separate populations of BA cells.

The following sections contain brief descriptions of the computational roles and interactions of these regions. A more in-depth description of our model architecture is provided in S1 Appendix. In particular, this includes model parameters defining three operational modes – *Perception* (memory formation), *Sleep* (memory consolidation), and

*Recall* (memory retrieval), whereas S2 Appendix breaks down the model’s update cycle.

### Hippocampal Formation (HIP & EC)

Our implementation of HIP draws on a computational model by Fiebig & Lansner [16]. During *Perception*, environmental inputs activate sparse subsets of HIP cells (4% active), and Hebbian learning strengthens synapses between co-active neurons to form a transient, context-specific *engram*. The overlap between two hippocampal engrams scales with the similarity of the contexts [25]. These representations are short-lived – in the sense that they rapidly become unlikely to entrap the network state [26] – as they are overwritten by new learning in HIP [18, 27–30].

During *Sleep*, in the absence of external inputs, recurrent excitation drives HIP to replay stored engrams. To prevent the indefinite replay of a single memory, inhibitory synapses between active cells are rapidly strengthened – using the same Hebbian rule on a fast timescale [16] – pushing activity toward other stored patterns. Though this implementation is oversimplified, neural inhibition does play a key role in terminating hippocampal replay events [31]. Multiple contexts are thus replayed sequentially over the course of *Sleep*. Plastic connections from HIP to both neocortex (CTX) and basal amygdala (BA_N_), formed during learning, enable coordinated sleep reactivation of context-specific engrams [6].

During *Recall*, HIP activity tends towards the engram of the ‘learned’ context that is most similar to the current environmental input. Input features of the current environment are compared to those associated with the activated HIP engram via a feedback circuit involving the entorhinal cortex (EC; Supplementary Fig. S1) [11, 32, 33]. If input and HIP engram match, the model attempts to retrieve an associated fear response via HIP*→*BA_N_ synapses. Otherwise (i.e., if HIP does not recognize the current context), the model instead engages CTX *→* BA_N_ synapses, relying on long-term cortical memory to determine if the current environment is associated with fear.

### Neocortex (CTX)

Our model implements the transfer and transformation of context memories from hippocampus to neocortex (*systems consolidation*) for long-term storage [34, 35]. Experimental evidence suggests that neocortical engram cells encoding a memory are selected during hippocampal memory formation, but initially lack sufficiently strong connections to be collectively reactivated by sensory cues alone [18]. In our framework, non-plastic synapses from SC to CTX determine the engram cells that go on to encode the current context during *Perception*, while plastic synapses from HIP to CTX are simultaneously strengthened. Therefore, when HIP replays context-specific engrams during *Sleep*, their CTX representations are reactivated and gradually consolidated via Hebbian plasticity. Simultaneously, slowly-evolving synapses from active CTX cells onto co-activated BA_N_ cells are strengthened. As a result, associations between context and fear responses initially supported by HIP→BA_N_ synapses progressively shift to stable CTX→BA_N_ synapses over time (cf. Fig 2; [18]).

**Fig 2.**
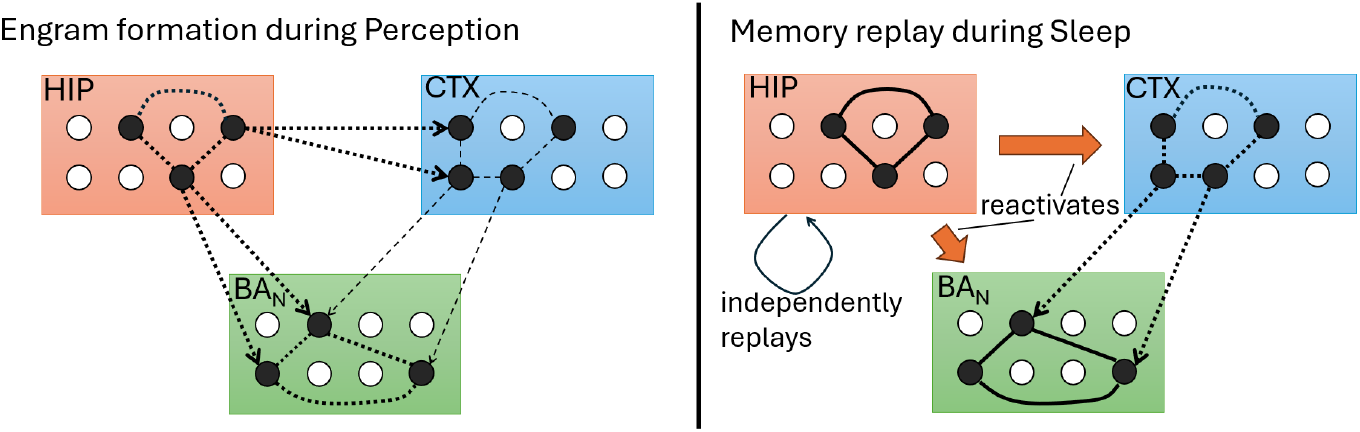
Dynamics of engram formation and replay. Sleep replay in HIP drives the consolidation of activity patterns in the CTX *→* BA_N_ circuit. Filled circles denote active engram cells encoding a specific context that is currently perceived (Left) or internally replayed (Right). Solid lines denote synapses that have formed in the past. Dotted lines denote synapses in the process of being strengthened. *Thin* dotted lines denote *slow* synaptic changes that generally do not suffice to form persistent associations. Hence, engram formation in CTX relies on sleep re-activations.

Due to its denser activation patterns, CTX engrams overlap more strongly than their HIP counterparts, capturing shared features of remotely experienced environments [36]. Our later results demonstrate that this decrease in memory separation entails increases in fear generalisation over time [37–40].

### Amygdala (BA & CeA)

Previous neural network models of context fear typically encoded the strength of fear associations directly in synapses connecting hippocampal engram cells to fear-promoting amygdala neurons [10, 12, 41]. A limitation of this approach is that transferring fear associations from hippocampus to cortex via replay becomes problematic: precise synaptic strengths would need to be copied or accurately reproduced across regions [17]. To address this, our model proposes that the magnitude of fear associations is instead encoded internally within the amygdala, specifically via context-specific amygdalar engram cells that recruit valence-coding neurons through intra-amygdalar synapses [18, 42]. Thus, coordinated sleep replay only needs to associate cortical context engrams with corresponding amygdala engrams – a simpler task (cf. Fig 2).

In our model, when a *prediction error* – a difference between the current aversive input (US) and the model’s fear response – occurs, recurrent synapses between active valence-neutral amygdala engram cells (BA_N_) are strengthened [43, 44], providing BA_N_ with a stable index of the current context. Depending on the sign of the prediction error, it further drives a strengthening of synapses extending from this BA_N_ engram onto separate populations of fear-promoting (BA_P_) or fear-inhibiting (BA_I_) cells [45, 46]. In this way, valence-coding neurons can persistently become associated with (‘recruited by’) a context. Active BA_P_ (or BA_I_) neurons drive the model’s fear response by exciting (or inhibiting) its single fear-output neuron, analogous to the central nucleus of the amygdala (‘CeA’) [47, 48].

### Recruitability

BA_P_ cells that have become associated with many different contexts may, as a result, receive strong excitation even in entirely novel environments – leading to indiscriminate fear expression. To prevent this scenario, our model limits synaptic plasticity during each learning episode to a small, time-varying subset of valence-coding neurons. This modulation is controlled by a cell-specific factor termed ‘recruitability’ (ℛ). During fear or safety learning, the prediction error gating plasticity onto each valence-coding neuron (in BA_P_ or BA_I_) is scaled by its current ℛ value. At any given time, ℛ is close to 1 for only a few cells, which are then more likely to undergo plasticity (Fig. 3a).

**Fig 3.**
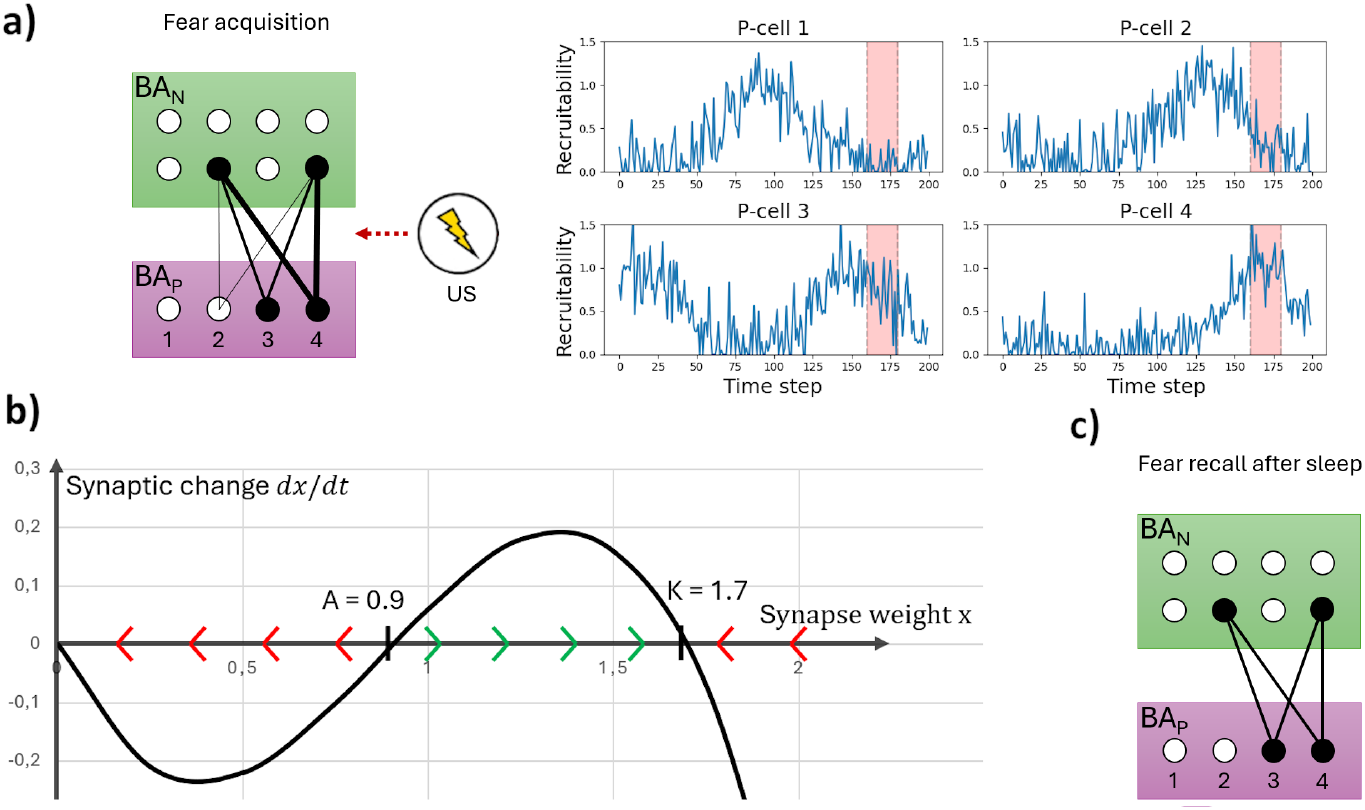
Fear acquisition and synaptic homeostasis. **a) Fear acquisition:** When a *surprising* US occurs, active (valence-**N**eutral) BA_N_ engram cells, encoding the current context, become associated with fear. **Left:** Over the course of several time steps, synapses from the BA_N_ engram onto fear-**P**romoting BA_P_ cells are strengthened by various amounts (denoted by the different line weights). **Right:** The extent to which synapses onto certain P-cells are strengthened is mediated by those cells’ current *recruitability* (Supplementary Fig S2). In this example, the time window of US delivery (time step 160 − 180) is shaded in red. P-cells 3 and (especially) 4, but not 1, are *recruited* – i.e., become associated with the current context – as the US signals coincide with their time windows of high recruitability. Synapses onto P-cell 2 are strengthened to some, limited extent as random noise has raised its recruitability for a single time step during conditioning. **b) Cubic growth model:** The differential equation describing homeostatic changes applied to the strength of our model’s valence-coding synapses during *Sleep*, 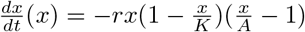 with (exaggerated) *learning rate r* = 1.4, *extinction threshold A* = 0.9 and *recruitment strength K* = 1.7. For *t → ∞*, the strength stably converges to *x* = 0 or *x* = *K*, depending on whether its starting value *x*(0) lies above or below *A*. This rule gives rise to the synaptic changes from **a)** (Left) to **c)**. **c) Fear recall after sleep:** When the conditioned context is encountered after a *Sleep* phase, synaptic homeostasis has acted on the synapses strengthened in **a)**. Weak synapses have been pruned, whereas stronger synapses have been normalized towards an appropriate strength. See also Fig 6 and its discussion in later results.

Our implementation of this mechanism (Supplementary Fig. S2; cf. S2 Appendix) was computationally motivated, and it does not reflect a specific biological signal. Nonetheless, the brain likely employs various processes to regulate neuronal allocation [49]. For example, neurons expressing higher levels of the transcription factor CREB exhibit increased excitability and preferentially become assigned to newly formed memory traces [50–52]. Our implementation reflects the finding that fear memories formed close in time to each other show greater neural overlap [51, 52].

### Homeostasis

The *synaptic homeostasis hypothesis* proposes that sleep globally weakens synapses to preserve functionally meaningful connections [53]. Experimental evidence confirms that synaptic homeostasis during sleep is selective, preferentially weakening synapses that are already weak, while relatively preserving stronger ones, as observed in sensory and motor cortices [54, 55]. Within the amygdala, sleep-dependent synaptic adjustments appear spine-type- and subregion-specific, though their precise functional implications remain unclear [56]. Nevertheless, these observations are intriguing given that sleep deprivation increases amygdala reactivity [57] and may facilitate future fear acquisition [58]. These findings raise the possibility that disruptions of synaptic sleep homeostasis promote amygdala hyperexcitability, contributing to maladaptive fear sensitization and generalisation – phenomena central to pathological anxiety [59].

In our model, synaptic homeostasis at connections onto valence-coding amygdala cells ensures adaptive fear learning. For valence to be stably associated with a context, synapses between context engram cells (BA_N_) and valence-coding cells (P-or I-cells) typically require repeated strengthening over multiple subsequent simulation steps during *Perception*. Although slow fluctuations in cell-specific recruitability promote the recruitment of specific neurons in specific conditioning events, random noise occasionally produces transiently high recruitability in some cells for a single time step (Fig 3a). This can produce partially strengthened, functionally ineffective synapses that do not reflect stable context-valence associations. Unregulated, such incidental synaptic changes could accumulate, resulting in inappropriate fear responses or accelerated acquisition (interpreted as non-associative fear sensitization [20]). Furthermore, *excessively* strong synapses could lead to inappropriate fear generalisation, as will become clear in later results.

To mitigate these issues, we implemented a synaptic homeostasis mechanism modelled via a *cubic growth function* (Fig. 3b). Although cubic growth has not previously been proposed as a biological model of synaptic change, our implementation reflects general experimental observations: during sleep, weak or incidentally strengthened synapses are preferentially pruned, while strong synapses are relatively preserved [54, 55] and may even be strengthened [60, 61] (Fig 3c).

## Results

In the following, we demonstrate how our model captures key mechanisms of fear memory formation, consolidation, and retrieval through targeted simulations. All simulation protocols are provided as supplementary materials in Tables S7 to S14.

### Context engram formation, replay and recall

In *Perception* mode, the activity patterns of our model’s engram modules HIP, CTX and BA_N_ depend on the features of the current environmental input. All three modules are capable of storing their activity patterns by strengthening excitatory synapses between co-active units, which thereafter form an *engram* that may be re-activated in *Recall* mode. Fig 4a illustrates the dynamics of *engram formation* in a context presented for 50 time steps. HIP swiftly formed a strongly connected engram, whereas CTX advanced much more slowly. BA_N_ was able to form an engram very rapidly, but only did so once a surprising US was delivered. At the same time, the model formed synapses from active HIP-to active CTX- and BA_N_ units, which would lay the foundation for coordinated replay across all three regions in a subsequent *Sleep* phase.

**Fig 4.**
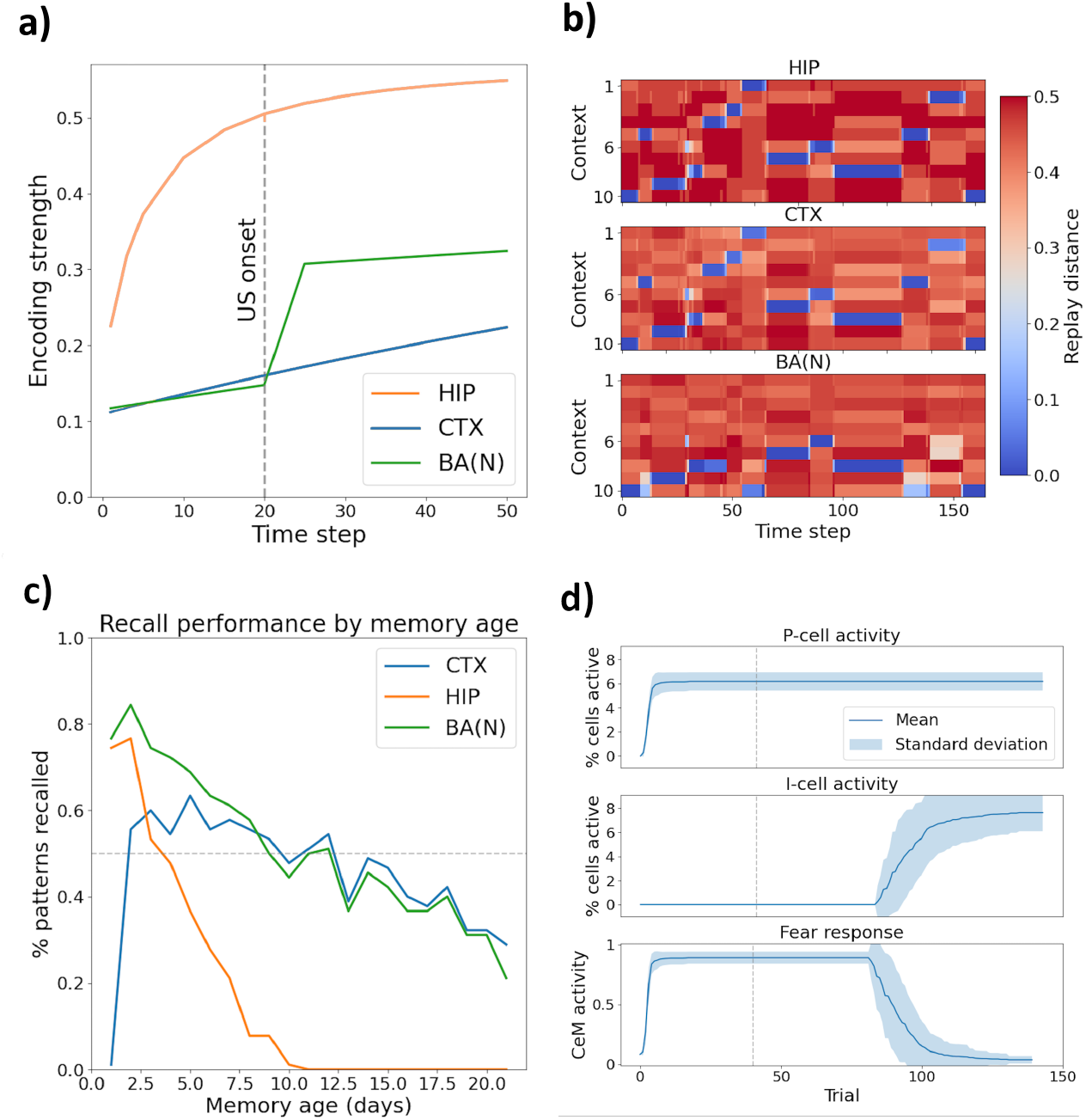
Mechanisms of context/fear memory formation. **a) Engram formation:** Encoding strength (net synaptic weight between active engram cells, normalized by their total outgoing weights) of a novel context, presented for 50 time steps, in HIP, CTX, and BA_N_. US inputs occurred on the last 30 time steps. HIP rapidly formed context engrams; BA_N_ did so after the first US; CTX learned slowly. Results averaged over 10 simulation runs (simulation protocol in Supp. Table S4). **b) Sleep replay:** Distance [64] between original context patterns and neural activity during *Sleep* in HIP, CTX, and BA_N_ following sequential exposure to 10 novel contexts. In contexts 6 to 10, US signals were delivered. Blue indicates replay of stored engrams. All contexts were replayed in HIP and CTX; BA_N_ replay was limited to contexts paired with aversive stimuli. Simulation details in Supp. Table S5. **c) Recall performance:** Fraction of contexts successfully recalled by HIP, CTX, and BA_N_ as a function of days since initial encoding. HIP reliably retrieved recently, but not remotely, perceived contexts. CTX was most likely to recall contexts encoded several days ago, reflecting sleep-dependent consolidation. Compared to HIP, CTX featured a recall curve with a lower peak, but flatter drop-off, indicating a more selective, more durable, storage strategy. BA_N_ recalled contexts when supported by either HIP or CTX (curves averaged over 40 runs; simulation protocol in Supp. Table S6). **d) Acquisition and extinction:** Fear acquisition and extinction dynamics. In each of 30 simulations, a context was presented for 140 steps, with US inputs delivered only during the first 40. The model acquired fear quickly (via BA_P_ activation), which was slowly extinguished following removal of the US through gradual activation of BA_I_ cells (Shaded regions = 1_SD_ across runs; protocol detailed in Supp. Table S7). BA_N_ *→* BA_I_ synapses began growing in strength as soon as the US signals were removed, but fear did not decrease until – after a considerable delay – the first BA_I_ cells became active.

Such sleep replay is illustrated in Fig 4b. This plot demonstrates the hippocampal replay of 10 contexts, previously shown to the model, over the course of a *Sleep* phase. Whenever HIP replayed a context, the corresponding CTX pattern was co-activated. BA_N_ participated in this coordinated replay only for contexts for which it had formed an engram – i.e., those which had been paired with US signals (contexts 6 to 10). Qualitatively, this aligns with the observed participation of the amygdala during the hippocampal replay of recent, threatening experiences in rats [9].

These replay dynamics, joined by Hebbian plasticity, allow synaptic connections to form between the CTX and BA_N_ engrams of emotionally salient contexts, allowing the latter to be recalled even after the corresponding HIP engram is forgotten. The recall performance of our three engram modules is demonstrated in Fig 4c. In this simulation, the model *perceived* three novel contexts on each of 25 days, intermitted by *Sleep* phases. Each context was joined by light US signals to engage plasticity in BA_N_. To assess the model’s ability to remember each context, it was ultimately placed in*Recall* mode. Each of the 75 original input patterns was ‘blurred’ (10% random masking) and presented to the model for one time step. If, after 30 time steps without further inputs, a module’s activity matched the original memory trace (within some margin of error, cf. Supp. Table S3), recall was ‘successful’. HIP consistently recalled recent contexts but failed to retrieve context memories older than approximately 10 days. In contrast, CTX’s ability to recall contexts initially increased over time, rising steeply after one night of sleep and peaking around five days post-encoding. The initial rise reflects ongoing consolidation of CTX engrams driven by hippocampal sleep replay. BA_N_, receiving inputs from both HIP and CTX, successfully recalled contexts as long as at least one of these regions had retained the associated memory.

In summary, replay of recently formed HIP memory traces during *Sleep* co-activates associated CTX and BA_N_ engrams. Hebbian plasticity strengthens the CTX engram’s synapses and links it to the co-activated BA_N_ trace. After one or several replay events for the same context, this mechanism allows the *Recall* of that context’s BA_N_ engram, when cued with a relevant input, even if the initial HIP encoding has grown too weak to be retrieved.

### Fear acquisition and extinction

For our model to explicitly associate a context with fear (or safety), it must form a BA_N_ engram, link it to the corresponding HIP trace and – downstream of BA_N_ – recruit valence-coding BA_P_ (or BA_I_) cells. Fig 4d shows a simple simulation of fear acquisition and extinction in the same environment. When a context was repeatedly paired with a US, the model rapidly developed a fear response through recruitment of fear-promoting BA_P_ cells. Following removal of the aversive stimulus, extinction occurred more gradually, as fear-inhibiting BA_I_ cells slowly became activated and suppressed the fear response. This mechanism for extinction – the recruitment of fear-inhibiting neurons downstream of the encoding of the fear association [45, 46, 62] – aligns with experimental findings suggesting fear extinction depends primarily on silencing, rather than erasure, of the original fear association [62, 63].

### Fear renewal and generalisation

In our model, fear (or safety) associations can generalize because contexts with shared input features tend to receive overlapping engrams in HIP, CTX and – in particular –

BA_N_. Hence, valence-coding cells linked with a conditioned context may become active when a similar environment is presented in *Perception* or *Recall* mode (cf. Fig 5a). Context-dependent fear renewal – the reappearance of previously extinguished fear upon encountering a context different from the extinction environment – is a common clinical challenge that undermines anxiety therapies [65]. It occurs because extinction memories (which suppress learned fear) tend to be more context-specific than the initial fear association [66].

**Fig 5.**
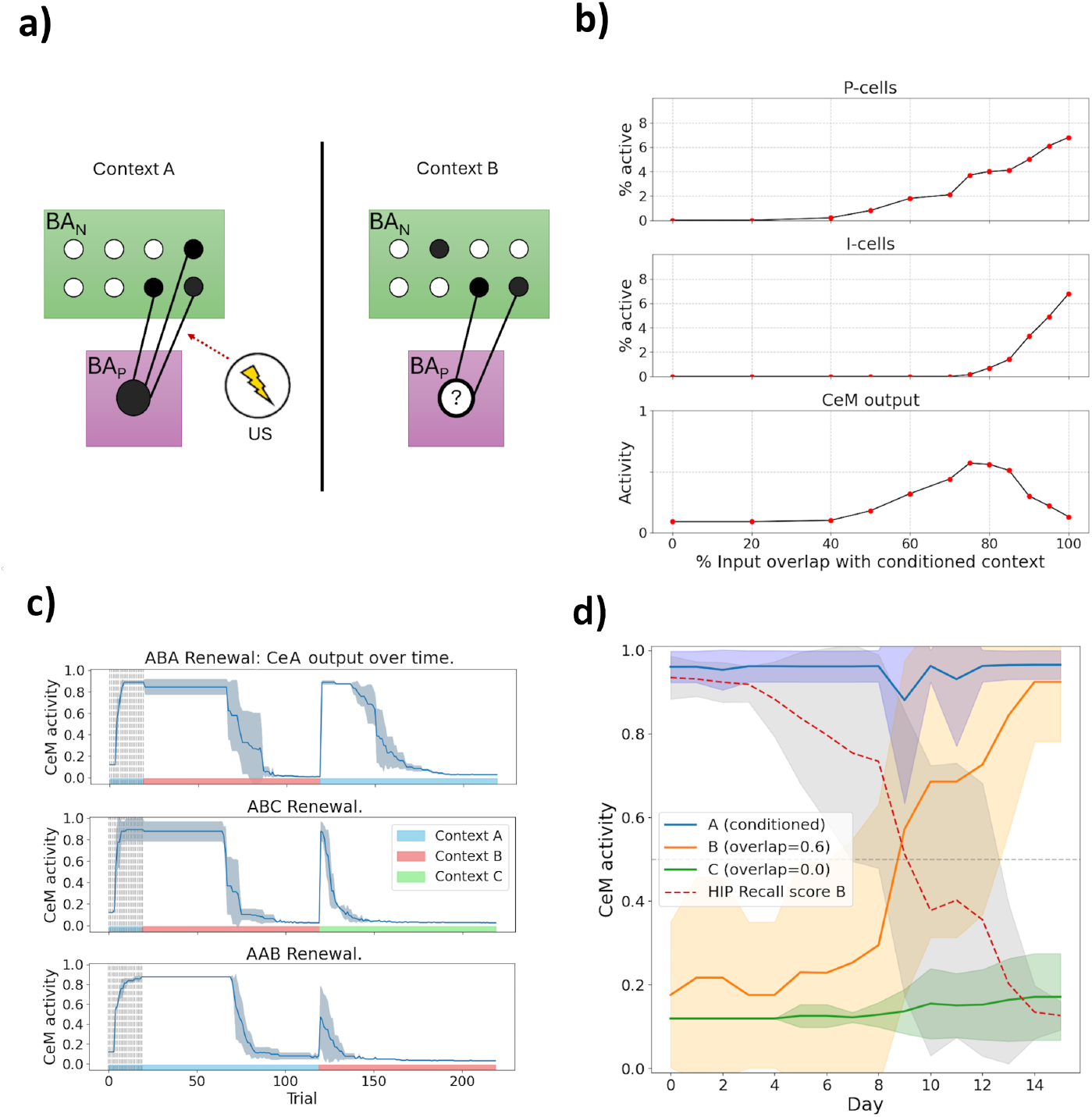
Results on fear generalisation. **a) generalisation schema**. Fear conditioning in context ‘A’ strengthens synapses between its *BA*_N_ engram and *BA*_P_ cells, activating the latter. In a similar context ‘B’, some of these *BA*_*N*_ cells are active, exciting the same *BA*_P_ cells. Whether these become active, causing fear to generalize, depends on the margin by which their activation threshold was exceeded in context ‘A’ and on the overlap of the two *BA*_N_ engrams. **b) generalisation gradients**. After fear acquisition and extinction in context ‘A’, red dots denotes the fraction of active BA_P_-cells (top), of active BA_I_-cells (center) or the magnitude of the CeA output (bottom) when the model meets a context whose feature overlap with ‘A’ is shown on the x-axis. Plots were averaged over 10 runs. Fear renewal is likeliest to occur when the model is placed in a context moderately similar to ‘A’. **c) Renewal paradigms**. From top to bottom, the figure shows the activity of our model’s CeA cell when subjected to the ABA, ABC and AAB fear renewal paradigms, as described in the main text. Plots were averaged over 10 runs. **d) Fear generalisation increases over time**. After fear conditioning in context ‘A’, fear expressed in a similar, unconditioned context ‘B’ increases as days pass. In a dissimilar context ‘C’, no fear is expressed, showing that the increase in ‘B’ is associative. The dotted ‘Recall score’ (F1 measure) quantifies how accurately the HIP engram retrieved when observing context ‘B’ in *Recall* encodes its true input features (Supplementary Fig S1) – it is inversely proportional to fear expression in ‘B’, indicating the generalisation increase is tied to use of the CTX *→* BA_N_ pathway for *Recall*. Curves were averaged over 30 runs. Shaded regions denote one standard deviation across runs.

We captured this asymmetry by assigning higher activation thresholds to fear-inhibiting (I-) cells compared to fear-promoting (P-) cells. In addition, our model’s Hebbian learning rules place an upper bound on the maximal strength of any synapse [27], which allowed us to choose parameters in a way ensuring no I-cell ever receives *much* more excitation than needed to become active. Consequently, only contexts highly similar to a conditioned environment activate enough of its BA_N_ cells to exceed I-cells’ thresholds, whereas P-cells become active even in moderately similar contexts, promoting broader fear generalisation.

To illustrate this, we trained the model on fear acquisition followed by extinction in a specific context (‘A’). Subsequently, we assessed the activation of P- and I-cells when the model encountered contexts with varying degrees of similarity to ‘A’ (Fig.5b). As intended, I-cells recruited during extinction required contexts to be highly similar (above 80% input overlap) to become reactivated. In contrast, P-cells could reactivate in contexts with as little as 50% input overlap with ‘A’, causing fear responses to generalize broadly. These dynamics led to realistic patterns of fear renewal across classical ABA, ABC, and AAB protocols, in which fear is first acquired (in context ‘A’), extinguished (in ‘A’ or ‘B’), and tested for renewal (in ‘A’, ‘B’, or a novel context ‘C’; Fig.5c). Notably, renewal magnitude followed the order *ABA > ABC > AAB*, in line with experimental observations [19, 67]. In all cases, extinction in renewal contexts occurred faster than initial extinction, reflecting sub-threshold excitation accumulated onto I-cells during prior extinction learning [68, 69].

Our model also captures increases in fear generalisation over time, consistent with experimental reports [37–40]. This emerges naturally through the transition from HIP- to CTX-dependent recall. Immediately after fear acquisition in context ‘A’, exposure to a moderately similar but harmless context ‘B’ initially does not elicit significant fear. However, as the context memories consolidate over time and retrieval increasingly relies on cortical representations - less sparse and more overlapping than their hippocampal counterparts [16], fear expressed upon *Recall* of context ‘B’ gradually increases (Fig. 5 d). This occurs because the cortical engram for context ‘A’ becomes partially activated when remotely recalling context ‘B’, thereby activating portions of the conditioned BA_N_ engram. Importantly, because BA_N_ forms specific engrams only for contexts linked with prediction errors during *Perception*, systems consolidation selectively increases fear expression in context ‘B’ without diminishing fear recall in ‘A’. Our model thus mechanistically explains the experimental observation that fear generalisation increases over time due to decreased memory specificity upstream of the amygdala—yet crucially, the original fear memory itself remains robustly recallable, consistent with empirical findings [39, 70, 71].

### Sleep homeostasis

#### Demonstrating the mechanism

Besides memory consolidation, sleep in our model plays a critical role in maintaining synaptic homeostasis within the fear circuitry. In any fear conditioning event (say, in context ‘A’), synapses from active BA_N_ cells onto fear-promoting (P-) cells are strengthened to varying degrees, modulated by each P-cell’s current *recruitability* (Fig 3a, Supplementary Fig S2). If the model later encounters a novel, similar context ‘B’, any P-cell receives a fraction of the excitation it did in context ‘A’ (Fig.6a,b). Even if the most strongly innervated P-cells remained below activation threshold, their subthreshold excitation would predispose the model to rapidly acquire fear in context ‘B’, should a mild aversive event occur.

While enhanced fear learning after stressful experiences is plausible [20, 21], unchecked, the above effect may eventually cause fear to be expressed in a broad range of environments reminiscent of context ‘A’. Addressing this is one role of our synaptic homeostasis mechanism (cf. ‘Methods: Model description’) – it depresses excessively strong synapses onto P-cells, thereby lowering the tendency of their recruitment to generalize. As illustrated in Fig. 6c-d, this homeostatic adjustment reduces the net innervation of ‘recruited’ P-cells in context ‘A’ towards a stable intermediate range, limiting associatively generalized fear in ‘B’. Additionally, the same homeostatic process also prunes weak synapses onto P-cells, to prevent a non-associative buildup of excitatory inputs over time – an effect we further examine in the following.

**Fig 6.**
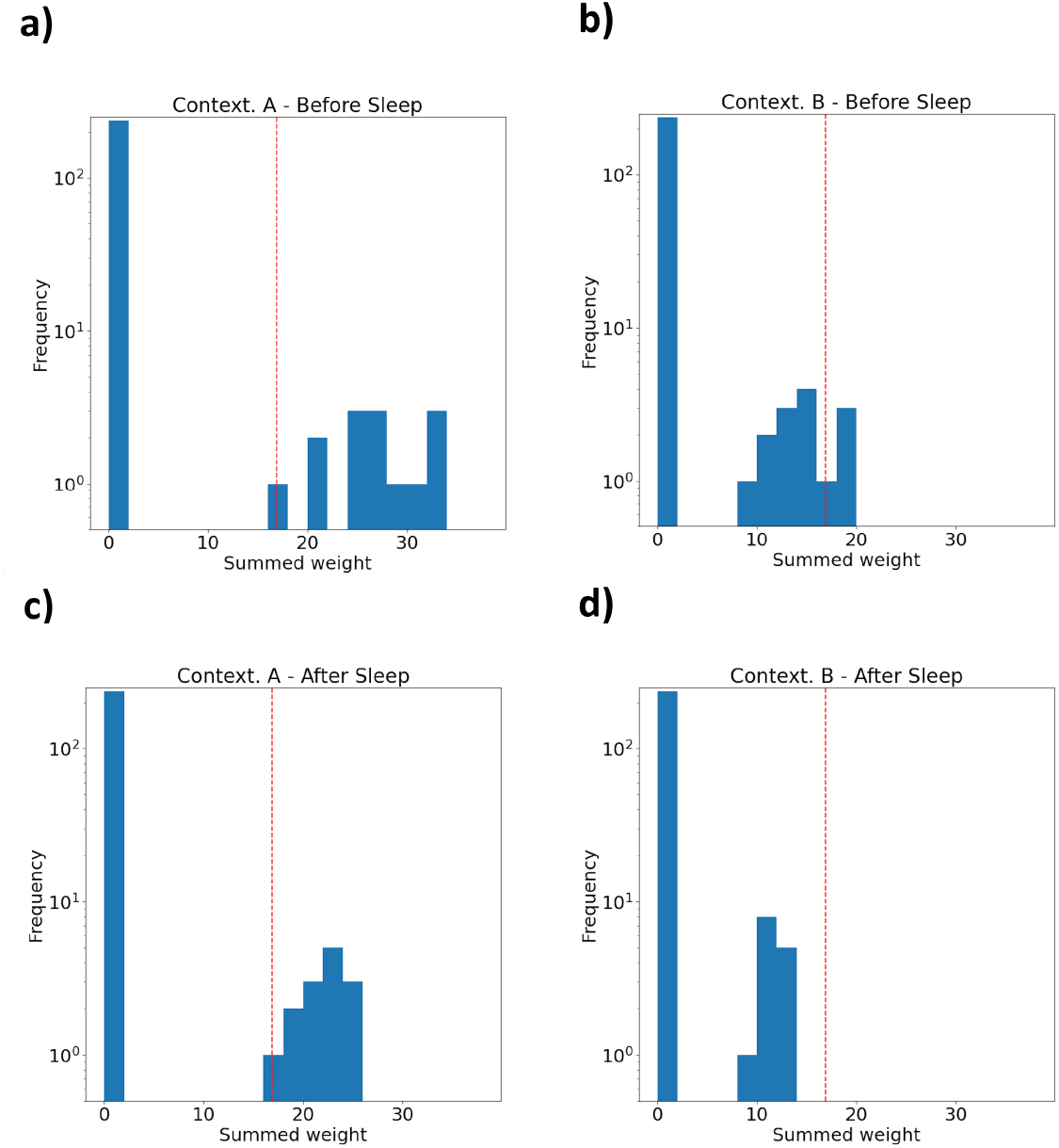
Histograms of the summed strength of afferent synapses from active BA_N_ units over all P-cells. The red lines symbolize the least amount of excitation a P-cell must receive to become active. Histograms were computed… **a)** …at the end of fear acquisition in context ‘A’. BA_P_ cells falling on the right of the red line are active, sparking fear. **b)**…upon being immediately placed in an unconditioned, vaguely similar context ‘B’. The BA_P_ cells that were most strongly innervated in ‘A’ are active – expressing some generalized fear. **c)**…upon revisiting context ‘A’ after a *Sleep* phase. The amount of fear expressed has not changed relative to a) but – thanks to homeostatic synaptic adjustments during *Sleep* – no BA_P_ cell receives much more excitation than needed to be activated. **d)**…upon revisiting context ‘B’ after a *Sleep* phase. Fear is no longer expressed in this unconditioned context. A selective normalization / partial reversal of recently strengthened fear-coding synapses in amygdala circuits during *Sleep* is thus posited to underlie decreases in the generalisation of freshly learned fear associations [80].

#### Sleep deprivation

Insomnia—a chronic difficulty in falling or staying asleep—is commonly observed in anxiety disorders [72, 73], and sleep disruptions may increase vulnerability to anxiety-related pathologies [74, 75]. Notably, sleep between fear conditioning sessions has been shown to counteract sensitization to experimental stressors [74, 76, 77].

Our model can explain these findings due to our assumption that sleep after fear learning prunes weak synapses onto P-cells. If such synapses are formed as a byproduct of fear conditioning events, or in response to negligible, moderately aversive events during daily experience, our model predicts that regular homeostatic pruning of unneeded *BA*_*N*_ *→ BA*_*P*_ synapses is essential. Without it, weak associations would accumulate, increasing the synaptic density of the amygdala fear circuit and enhancing fear acquisition and expression.

To demonstrate this prediction, we designed a simulation that mimics daily experiences alongside sleep-induced synaptic homeostasis. Over seven *days*, the model encountered five novel contexts each day, each paired with moderate US signals. Between days, the model underwent a *Sleep* phase. On the eighth day, the model was briefly exposed to a novel context with a moderate US (strength 0.5) for five time steps, after which its fear response was recorded. Repeating this protocol while systematically varying the duration of the *Sleep* phases revealed that shortened sleep significantly accelerated fear acquisition in the novel context (Fig 7a) and led to net increases in *BA*_*N*_ *→ BA*_*P*_ synaptic weights (Fig 7b).

**Fig 7.**
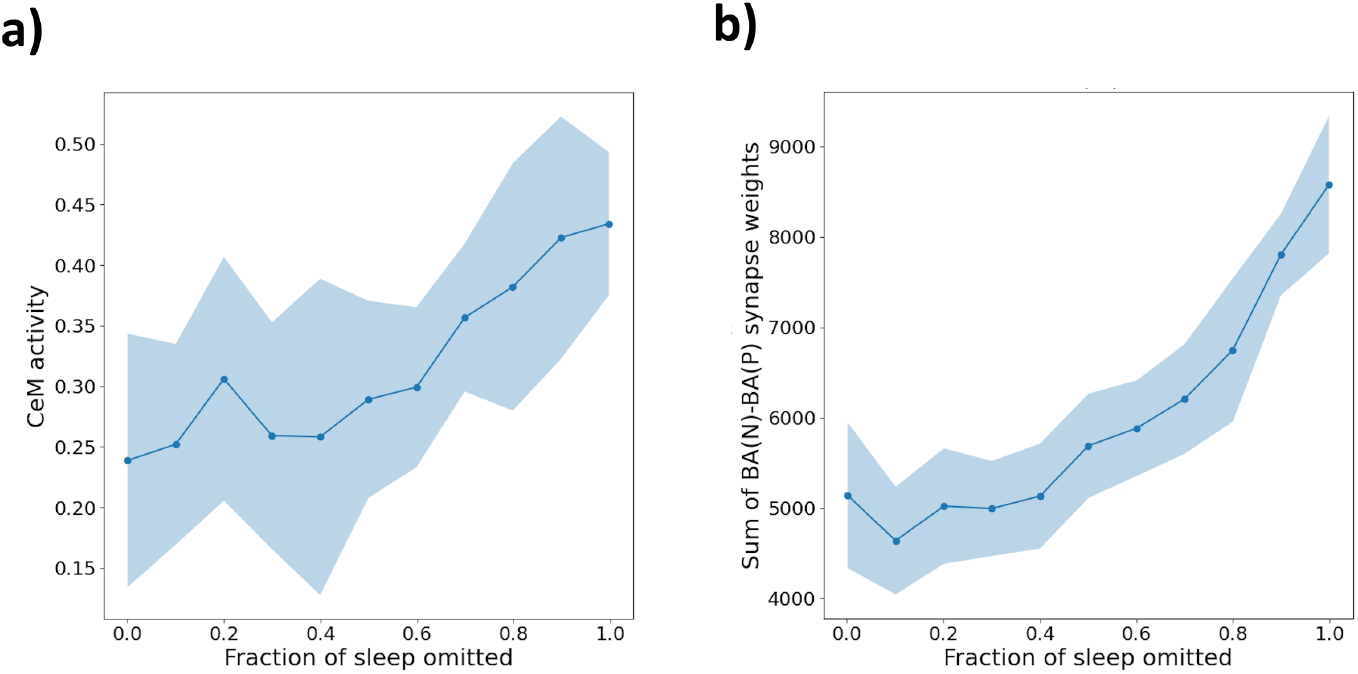
Effects of chronic sleep deprivation. Instances of the model were subjected to 7 days filled with exposure to various contexts, joined by moderate US signals. At the end of the simulation, fear acquired after the delivery of 5 moderate US signals in a novel environment was tested. **a)** The magnitude of the fear response learned in the novel environment, at the end of the above protocol, increased as the daily duration of *Sleep* was systematically lowered from a default of 165 time steps. **b)** Sleep deprivation was further accompanied by net increases in the strength of synapses from context-onto fear-coding BA cells, measured at the end of the simulation protocol. In both plots, dots denote the mean, shaded regions one standard deviation across 10 runs of the simulation.

These results suggest that regular sleep-induced synaptic pruning is critical for mitigating non-associative fear sensitization. Consequently, targeting sleep disruptions —an approach that has, e.g., already shown promise in reducing PTSD symptoms [78, 79]—may directly influence the synaptic mechanisms underlying chronic anxiety. Our model thus makes the testable prediction that enhancing synaptic sleep homeostasis in the amygdala could be a key therapeutic strategy in PTSD and related disorders.

#### Stress-Enhanced Fear Learning

In the previous section, we demonstrated that sleep deprivation disrupts synaptic homeostasis in the amygdala, allowing weak synapses to accumulate, thereby enhancing fear sensitization. However, impaired synaptic pruning may not be limited to disrupted sleep alone. Psychological stress also affects emotional reactivity and is associated with lasting synaptic potentiation and increased excitability in the amygdala [81–83]. Given these parallels, we hypothesize that severe stress can similarly disrupt synaptic downscaling processes during sleep.

Stress-Enhanced Fear Learning (SEFL) describes the phenomenon whereby exposure to severe stress in one context (‘A’) subsequently enhances fear acquisition in an unrelated context (‘B’) compared to non-stressed controls [20]. Notably, enhanced fear occurs specifically when stress precedes—but not when it follows—mild fear conditioning [20], indicating a primarily non-associative fear enhancement effect rather than associative fear generalisation [21]. Experimentally, extreme acute or chronic stress reliably induces lasting increases in long-term potentiation (LTP) within the basolateral amygdala (BLA), potentially priming neural circuits for heightened fear learning and generalized fear responses [81–83].

We modelled such stress-induced disruptions by temporarily increasing the *recruitability* of fear-promoting (P-) cells, thus enhancing their likelihood of forming lasting associations with BA_N_ context cells. Additionally, we lowered the *extinction threshold* of the homeostasis rule acting on BA_N_ *→* BA_P_ synapses, similarly favoring synaptic strengthening over pruning during *Sleep*. These parameter changes were activated only by prolonged exposure to sufficiently intense aversive events (cf. S2 Appendix), conditions that did not occur in our previous simulations, making the present results complementary and independent. After the cessation of extreme stress, parameters gradually returned to their default values over the course of two to three simulated days.

We tested this implementation using a protocol designed to capture key elements of the SEFL paradigm [20, 21]. First, the model was exposed to extreme stress (high-intensity US) in context ‘A’, followed by a *Sleep* phase. On the subsequent day, the model underwent mild fear conditioning (brief, moderate-intensity US) in a novel context ‘B’, again followed by *Sleep*. Finally, fear recall in context ‘B’ was measured. Consistent with experimental observations, this protocol resulted in enhanced fear expression in context ‘B’ compared to unstressed controls (Fig.8a). Importantly, the SEFL effect did not occur when extreme stress followed moderate fear conditioning (Fig.8b), matching empirical findings [20]. Furthermore, the enhancement of fear acquisition was persistent, which we assessed by interposing 15 *Perception-Sleep* cycles involving mildly aversive experiences between conditioning in contexts ‘A’ and ‘B’ (Fig. 8c). Previously stressed animals still acquired more fear in context ‘B’ because, although the described increases in P-cell recruitability had vanished by the time of conditioning, the associated net increases in synaptic strength between *BA*_N_ and *BA*_P_ persisted throughout the simulation (Fig. 8d). This result aligns with the experimentally observed long-lasting impact of traumatic stressors in rats [40].

**Fig 8.**
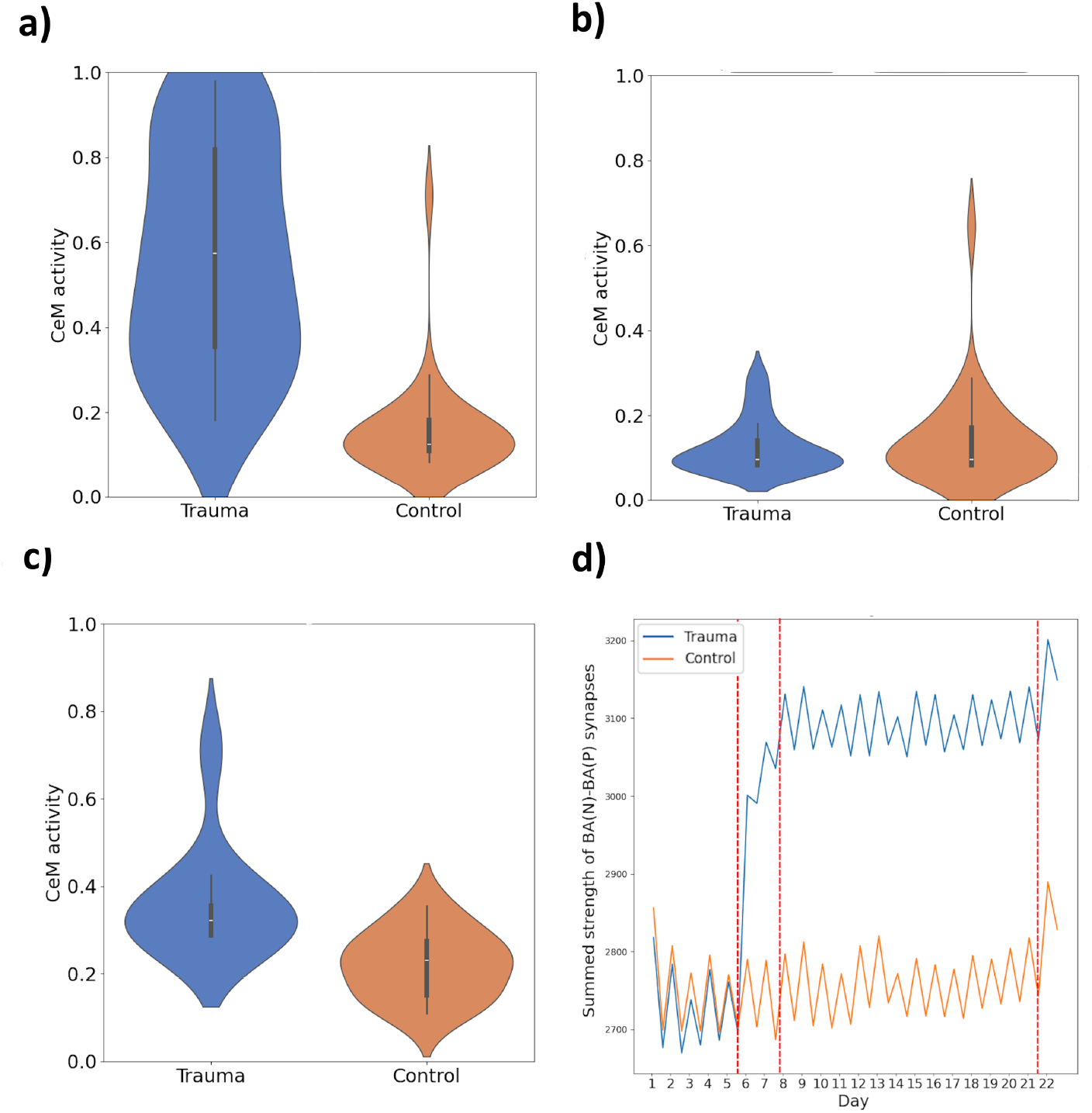
Results on stress effects. **a) Stress-Enhanced Fear Learning (SEFL)**. Violin plots describe the magnitude of the model’s fear response to context ‘B’, following the SEFL protocol as described in the main text, across 30 runs – with exposure to ‘traumatic stress’ on the one hand and without (‘control’) on the other. The trauma group acquires stronger fear responses to context ‘B’, showing fear sensitization. **b) SEFL does not occur if fear learning precedes stress**. Same as a), but with the order of ‘traumatic stressor’ and ‘moderate US’ exposure reversed. Trauma exposure does not retrospectively enhance fear of context ‘B’. **c) Stress enhances fear learning weeks later**. Same as a), but with ‘moderate fear acquisition’ in context B carried out 15 *Perception*-*Sleep* cycles after traumatic stress. During the delay, the model meets various contexts (with various, generally low threat levels). Fear is enhanced post-trauma. **d) Stress raises the net strength of** *BA*_*N*_ *→ BA*_*P*_ **synapses**. Blue and orange lines show the total, summed strength of *BA*_*N*_ *→ BA*_*P*_ synapses of the ‘trauma’ and ‘control’ model instances, averaged across the 30 runs that gave rise to c). Red vertical lines denote, from left to right, delivery of traumatic stress, the time by which the ‘trauma’ model’s homeostasis rule fully recovers and delivery of the ‘moderate US’. Values were recorded at the end of each *Perception* and *Sleep* phase, *explaining the oscillations* of the curves. *Sleep* entails net decreases in synaptic strength [84]. Note that those ‘falls’ are smaller while homeostasis is impaired, contributing to the trauma’s long-term impact.

Thus, our model demonstrates a unified principle: impaired homeostatic pruning of amygdala synapses — whether due to sleep deprivation or severe psychological stress — may predispose neural circuits to maladaptive fear sensitization. Clinically, this hypothesis – if validated – implies that therapeutic interventions targeting either sleep quality or resilience to stress-induced homeostatic dysregulation could reduce vulnerability to pathological fear and anxiety disorders.

## Discussion

In this paper, we presented a biologically inspired neural network model of associative fear memory formation explicitly incorporating sleep-dependent consolidation processes. Positing coordinated replay of context-coding cell populations in the hippocampus, amygdala, and neocortex, our simulations provided a mechanistic account for the observed transfer of context-fear memories from hippocampal-amygdalar to amygdalo-cortical circuits. Assuming cortical engrams overlap more strongly than their hippocampal counterparts, our model naturally reproduced empirically observed increases in fear generalisation following contextual fear conditioning. We found that the summed synaptic strength between context- and fear-coding cells provided a heuristic measure of the model’s level of ‘fear sensitization’—a largely context-independent enhancement of fear acquisition and expression. Drawing on the synaptic homeostasis hypothesis of sleep, we concluded that impaired synaptic weakening in amygdalar fear circuits – as might plausibly occur due to sleep disruptions or psychological stress – could contribute mechanistically to such sensitization. Although the simulations described in this paper were not intended to capture the complete pathophysiology of any specific disorder, our broader motivation for explicitly including sleep-dependent processes in a fear-learning model was to narrow the theoretical gap between traditional associative fear models and the gradual, often delayed, emergence of pathological anxiety observed in disorders such as PTSD. In the following, we discuss specific experimental predictions arising from our model’s conception of sleep’s role in fear memory consolidation, as well as experimental evidence supporting predicted links between disrupted sleep-dependent homeostasis, stress-induced synaptic changes, and enhanced fear learning. We conclude by highlighting potential implications of these findings for understanding the pathophysiology of anxiety disorders characterized by generalized fear.

### Sleep replay and circuits for fear memory consolidation

In our model, hippocampal pattern replay during *Sleep* supports the consolidation of context-fear associations by simultaneously reactivating cortical and amygdalar cell ensembles. These replay events promote the maturation of persistent cortical engrams and allow these engram cells to form robust connections with downstream amygdala neurons. Although precise timing and dynamics remain under investigation, this general hypothesis aligns well with recent empirical studies, notably those of Kitamura et al. (2017) [18].

Kitamura et al.’s findings—corroborated by recent literature—highlight key features that inform our model:

1. In rats, contextual fear conditioning (CFC) activates groups of ‘engram’ cells in the hippocampus, medial prefrontal cortex (mPFC), and basolateral amygdala (BLA). Hippocampal engrams initially support fear memory retrieval but become functionally silent within two weeks [8, 18]. In contrast, mPFC engrams—active during memory formation—transition from an initially silent state to being essential for remote recall, consistent with the role of CTX in our model [18, 85–87]. BLA neurons remain essential for fear recall at all stages; while many neurons participate broadly across multiple fear memories [88], each memory recruits a small subset forming stable, memory-specific engrams consistently reactivated during recall [18, 89].
2. Long-term retention of contextual fear memories depends on the maturation of their mPFC engram, involving a strengthening of recurrent synapses between initially silent cortical engram neurons [18, 85]. This synaptic maturation depends on sustained hippocampal inputs following learning; disrupting hippocampal-to-cortical communication impairs cortical engram consolidation and long-term memory expression [18, 90]. Hippocampal sharp-wave ripples (SPW-Rs)—key neural events implicated in hippocampal-cortical replay, particularly during sleep [91]—are essential for the long-term retention of contextual fear memories [92–94].
3. Remote recall of contextual fear memories increasingly depends on projections from the mature mPFC engram to the BLA. Experimental manipulations indicate that mPFC-to-BLA projections become essential for memory expression approximately two weeks after CFC in rats [18]. Consistent with this, recent experimental findings by Lee et al. (2023) suggest a gradual strengthening of synapses between mPFC and BLA engram cells encoding a specific context-fear association over the course of a month following its formation [85]. Selective strengthening of cortical-to-amygdalar synapses has also been observed in the hours following fear conditioning [95], possibly primed by postsynaptic increases in AMPA receptor function within the prelimbic-to-BLA pathway [96].

It is important to note that the temporal dynamics described above are specific to CFC. In contrast, the recall of fear associated with an elementary cue, such as an auditory tone, is hippocampus-independent even when the memory is recent, and engages a multisynaptic cortical-to-amygdalar pathway involving the paraventricular thalamus (PVT) [97, 98].

In short, the above findings suggest that the neural encodings of learned context-fear associations undergo *systems consolidation* – a shift from hippocampal to neocortical dependence for recall – as they age [99]. This is worth pointing out, as systems consolidation is usually considered within the context of declarative, or episodic, memories – which involve the explicit recollection of an event and do not generally include contextual fear memories [100, 101].

Consistent with our model, it has previously been proposed that, to retain the functional identity of the initial hippocampal memory trace, cortical engrams must acquire synaptic connections onto downstream, subcortical regions that resemble those of the original trace. This process is understood as involving the ‘replay and synchronization of learning-associated neuronal activity patterns in hippocampus and cortical regions’ [99]. However, it remains unclear by what mechanism the cortical engram acquires synaptic connections onto subcortical, such as amygdalar, memory engram cells. Our model demonstrates that, if hippocampal replay of an activity pattern associated with a CFC learning event co-activates corresponding cortical and amygdalar engram cells, this may recruit Hebbian plasticity to forge direct connections between them – providing a biologically plausible route for the cortical engram to ‘inherit’ the output pathway necessary for fear expression.

While this proposal aligns well with the reviewed literature, it is important to note that near-simultaneous re-activations of cortical and amygdalar engram cells during sleep have not yet been demonstrated – although broader findings on coordinated activity across hippocampus, mPFC and amygdala following fear learning support their plausibility [9, 102].

In summary, our model offers a mechanistic account of how cortical fear engrams may acquire functional connectivity with subcortical targets, consistent with theoretical proposals and indirectly supported by experimental findings. We specifically assume the existence of amygdalar engram cells that serve as stable ‘anchors’ throughout systems consolidation—initially targeted by hippocampal inputs, and later by neocortical ones such as from the mPFC.

In our model, these BA_N_ cells form a recurrently connected memory index, receive converging hippocampal and cortical inputs with a sleep-dependent shift in dominance, and project to downstream valence-coding neurons. Hebbian plasticity recruited during sleep co-activation of mPFC and BLA engram cells enables the cortical engram to inherit the output pathway originally established by the hippocampus. Here we posit that this occurs during hippocampal replay events, though co-activations of mPFC and BLA engram cells at other time points may serve the same purpose.

A key test of this mechanism would involve identifying the post-learning window during which mPFC–BLA engram co-activation emerges, then specifically disrupting synaptic plasticity within this period. Our model predicts that this would selectively reduce defensive responding at remote, but not recent, time points—without preventing recall of the conditioning context itself.

### Sleep homeostasis, stress and fear sensitization

Many existing neural network models of contextual fear conditioning (CFC) rely on synapses connecting context-coding neurons, activated by environmental inputs, to valence-coding neurons driving defensive responses [10, 11, 17, 41, 103]. Although assuming context- and valence-coding neurons form distinct, well-defined populations is likely an oversimplification, this abstraction provides a valuable conceptual framework for understanding how fear associations are encoded, stored, and retrieved.

A central tenet of the *synaptic homeostasis theory* of sleep is that synaptic strengths accumulated during waking experiences undergo global down-regulation during subsequent sleep periods [53]. Evidence from cortical circuits suggests that this down-regulation preferentially prunes weak synapses while preserving stronger, functionally significant connections—potentially normalizing their strength rather than eliminating them entirely [54, 55]. Our model investigates how applying these homeostatic principles specifically to fear-related synapses within a neural network framework influences fear learning and expression, yielding two key insights:

### Normalizing strong synapses prevents excessive fear generalisation

In our simulations, synapses linking context-coding cells to fear-coding neurons form with varying strengths influenced by existing synaptic weights, cell-intrinsic recruitability, and the intensity of aversive experiences (cf. Fig. 6a). Within the framework laid out in our results, the strongest of these synapses can promote maladaptive fear (over)generalisation. Our model predicts that sleep-dependent synaptic homeostasis counteracts this by partially weakening overly strong connections, thus maintaining appropriate context specificity of fear expression. Fear expressed in a conditioned context may yet increase after sleep, as our homeostasis rule allows a strengthening of synapses onto ‘nearly recruited’ fear neurons (BA_P_ cells).

### Regular pruning prevents build-up of synaptic strengths

Our simulations further predict that regular pruning of weak, functionally insignificant synapses in amygdalar fear circuits – putatively formed as a by-product of fear learning, or benign daily experiences – is critical to avoid cumulative increases in net synaptic strength. Without this pruning, fear-coding neurons would receive a gradually increasing amount of ‘baseline excitation’ in any unconditioned context. Consequently, new fear associations may form more rapidly and be expressed more strongly, even in the presence of merely moderate threat signals.

Sleep deprivation—whether acute or chronic—increases amygdala reactivity to negative emotional stimuli [57, 104–106]. In humans, this effect is associated with emotional hyperreactivity and is commonly attributed to impaired top-down regulation by the mPFC. Consistent with this, sleep deprivation *prior to* fear conditioning experiments has been linked with increased fear expression during training and accelerated fear acquisition [107, 108]. We hypothesize that sleep regulates future emotional reactivity, including fear expression, by decreasing synaptic strengths in excitatory input pathways onto amygdala cells responsive to aversive stimuli.

Rodent, as well as human, studies further support the prediction that sleep omission in the immediate aftermath of – cued or contextual – fear conditioning impairs the behavioural discrimination between conditioned and safe stimuli [80, 109–111]. In our model, as per Fig 6, the beneficial effect of sleep in this regard relies on a normalization of synapses implicated in recent fear learning events – which may strengthen or weaken individual synapses – consolidating learned associations to support robust recall, while limiting maladaptive generalisation, as described.

A testable hypothesis that can be derived from our model, in conjunction with the theory that low levels of certain neuromodulators (such as noradrenaline, serotonin and histamine) during sleep underlie its role of promoting synaptic weakening [53], is that augmenting these neuromodulators during sleep should reproduce effects similar to sleep deprivation, enhancing subsequent fear acquisition or reducing the specificity of learned associations.

Notably, psychological stress is known to enhance the release of noradrenaline and has been shown to promote long-term potentiation (LTP) relative to long-term depression (LTD) within the amygdala [81, 112, 113]. In line with the above, chronic stress in the days and weeks before fear conditioning experiments has indeed been linked with heightened excitability in amygdala circuits and enhanced fear acquisition [114, 115]. Although acute stress caused by ongoing experiences similarly increases amygdala excitability, suggesting a direct influence of the noradrenergic system on emotional reactivity [116], our model further predicts that stress drives persistent plastic changes in amygdala fear circuits by disrupting synaptic weakening that should occur during sleep. Within our framework, synaptic net increases in pathways providing excitation to fear-coding neurons may manifest as a form of context-independent fear sensitization during future fear learning. They can be long-lasting as, once consolidated, synapses whose strength has accumulated under stress would not be homeostatically pruned once stress ceases.

Following this line of reasoning, our model yields the hypothesis that psychological stress impairs fear regulation specifically by disrupting sleep-dependent synaptic weakening in amygdala fear circuits. If experimentally confirmed, this mechanism could explain how acute stress can persistently alter fear expression and learning, potentially clarifying one pathway by which stress exposure increases vulnerability to anxiety or trauma-related disorders in susceptible individuals.

## Conclusion and Future Directions

Here we have presented a neural network model for the formation and consolidation of associative context-fear memories. The model proposes that maturation of a cortical-amygdalar engram supporting remote recall of contextual fear memories [18] relies on simultaneous co-activation of cortical and amygdalar engram cells, driven by hippocampal replay events during sleep.

We further hypothesize a synaptic homeostasis mechanism within the amygdala, in which sleep regularly prunes weak and functionally insignificant synapses. Such pruning could counteract the gradual accumulation of synaptic weights, preventing the emergence of a chronic fear-sensitized state. Stress is proposed to disrupt this homeostatic mechanism, leading to persistent increases in amygdalar synaptic density and priming the brain for enhanced fear expression and acquisition. Experimentally testing this hypothesized causal pathway could yield clinically relevant insights. Heightened fear sensitization, excessive generalisation, and inflexible emotional learning—potentially resulting from saturation of synaptic plasticity [117]—are hallmark symptoms of PTSD [83, 118, 119]. These symptoms frequently co-occur with chronic sleep disruptions, particularly in REM sleep [74]. Our proposal aligns with evidence of locus coeruleus (LC) hyperactivity in PTSD [120], potentially mediating hyperresponsiveness to threatening stimuli [121]. LC projections to the amygdala become hyperactive following stress [122], whereas their silence during *healthy* REM sleep likely facilitates critical synaptic weakening processes [84, 123].

By explicitly incorporating sleep-dependent consolidation processes, our model thus identifies a clinically relevant pathway—linking psychological stress, disrupted synaptic homeostasis during sleep, and subsequent emotional sensitization—that extends beyond existing computational models of fear learning. This conceptual framework highlights qualitative relationships among stress exposure, sleep dynamics, and fear regulation, offering clear hypotheses for experimental validation.

Given its broad scope, the current model implements several processes—including synaptic homeostasis, stress-induced plasticity changes, and neuronal allocation—at a relatively high level of abstraction. Future computational studies can leverage this qualitative framework to explore specific biological mechanisms more deeply, including neuromodulatory dynamics (e.g., noradrenergic modulation via the LC), cellular-level plasticity mechanisms, and molecular pathways underlying synaptic homeostasis.

Addressing these detailed mechanisms is critical, as the neural underpinnings of fear memory processing and their dysfunction in neuropsychiatric disorders remain sparsely understood. Increasing biological specificity in computational models will facilitate the generation of experimentally testable predictions, enhance interpretation of empirical findings, and support the development of targeted interventions. Pursuing these directions is essential for bridging computational neuroscience with clinical advances in treating fear-related disorders.

## Data availability statement

All code written in support of this publication is publicly available on Zenodo (DOI: 10.5281/zenodo.15546751).

## Acknowledgements

This work was supported by the United Kingdom Research and Innovation (grant EP/S02431X/1), UKRI Centre for Doctoral Training in Biomedical AI at the University of Edinburgh, School of Informatics.

## S1 Appendix Supporting Information Formal Model Description

As per Figure 1, each *module* of our model, such as HIP, CTX or BA_N_, belongs to one of three kinds of network – Bayesian Confidence Propagation Network (BCPNN), k-Winner Takes All (kWTA) or ‘Binary’. We adopted the auto-associative BCPNN architecture from Fiebig et al. [16]. A full discussion of the network architecture can be found there, or in Sandberg et al. [27]. Below, we provide a full formal description of our model. But first, we provide, as a summary, a broad description of the technical building blocks on which our implementation is based.

CTX, our model’s ‘long-term memory module’, is a Bayesian Confidence Propagation Neural Network (BCPNN). This network archetype, developed by Lansner and colleagues [124], was shown capable of acting as an auto-associative ([125]) memory system [16, 27]. Its units are grouped into ‘hypercolumns’, within which they weakly compete for activity – a principle observed in the cerebral cortex [126]. The network is fully connected, with recurrent weights *W* enabling excitatory inputs, whose strength is modulated by a ‘gain’ or ‘conductance’ *g*_*L*_. Weights are updated via a Hebb-like learning rule, derived from a Naive Bayes classifier [27], at a rate inverse to time constant *τ*_*L*_. Optionally, to prevent the indefinite repetition of an attractor state in the absence of further input, complementary weights *V*, updated using a much lower (faster) time constant *τ*_*A*_, may inhibit excitatory inputs, mediated by *negative* gain *g*_*A*_. In the biological brain, a corresponding mechanism for discouraging neural patterns that have recently fired is observed, e.g., as the temporary depression of synapses between pyramidal cells recently involved in the same ‘population burst’ [127]. The updating equations defining a BCPNN are outlined in Appendix S1 Appendix and discussed in greater detail by other sources [16, 27].

In a BCPNN, a softmax function is used on any time step to determine the neurons’ activities within each hypercolumn (cf. S1 Appendix). The fraction of strongly contributing units may vary considerably between different attractors. This feature is one reason we – like Fiebig & Lansner before us [16] – opt to use this architecture only for our model’s cortical module. In subcortical regions of the biological brain that form engrams to encode sensory or contextual inputs, the size of these engrams is remarkably consistent (with a sparsity of, e.g. 2 − 6% in dentate gyrus or 10 − 20% in lateral amygdala [128]). To enforce such stable engram sizes, our model’s ‘engram layers’ apart from CTX use a kWTA-rather than softmax rule to compute neurons’ activities from their inputs – only the *K* = *α * N* (sparsity times network size) units with the highest net input are active at any point in time. We thus distinguish between BCPNN-(CTX) and kWTA (HIP, BLA) modules. However, the Hebbian learning rules for updating the modules’ weights (*W* and *V*)-apply to both kinds equally.

While this kWTA approach is suitable for storing a fixed-size engram – a ‘discrete index’ – for a given input, it does not allow the corresponding network to become inactive without further modulation. To associate a continuous quantity with a given engram, past models have assumed the needed information to be encoded in the (real-valued) strength of synapses extending from the engram cells [11, 17, 41].

Demonstrating an alternative mechanism, we allow the engrams of our model to recruit individual (e.g., fear-coding) cells out of a larger population to encode an associated quantity in the number of such neurons that are activated. For simplicity, we model such populations as sets of units without recurrent synapses. Any unit receives its input solely from synapses from other modules (see below). If the net input exceeds a chosen threshold, the neuron (which we then call ‘recruited’) fires (at a rate of 1).

Lastly, any two modules may be uni-directionally linked by a *Connection* instance.

The input from source unit *i* to target unit *j* is computed by multiplying current activity *π*_*source*,*i*_ with *g*_*F B*_ *\* log*(*W*_*ij*_), where ‘feedback gain’ *g*_*F B*_ is predetermined for each *Connection* and where the updating rules for weights *W* follow naturally from the BCPNN architecture. Alternatively, a *Connection* may be non-plastic, in which case *W* is set or randomly generated when the model is initialized.

In the following, our architecture’s building blocks are outlined in more detail.

### BCPNN

Here we present the core updating equations of the BCPNN. The model operates in discrete time steps of length *dt* = 10*ms* (though the model should not be interpreted as strictly operating on this timescale; rather, its dynamics qualitatively represent repetitive neural firing over longer durations). On each time step, the net excitatory input *h* received by each unit in a BCPNN evolves according to Equations (1) through (10) below.

The network is organized into hypercolumns *H*_*k*_, each containing *M* units. The activity of units within a hypercolumn is normalized, simulating lateral inhibition mediated by inhibitory interneurons. This organization allows the network to operate as a soft-winner-takes-all (WTA) system. Groups of neurons that qualitatively operate in this manner have been described as the ‘basic functional units’ of neural computation in the cerebral cortex [126].

### Dynamics of Net Input

The net input *h*_*j*_(*t*) received by unit *j* is updated based on the activity of other units in the network and the synaptic weights. Two types of connections contribute to this update:

- **Excitatory connections**, with weights *w*_*ij*_(*t*) and biases *β*_*i*_(*t*), which implement Hebbian learning.
- **Inhibitory connections**, with weights *v*_*ij*_(*t*) and biases *γ*_*i*_(*t*), which implement an adaptation mechanism that discourages prolonged firing of the same cell ensemble.

The net input evolves as follows:

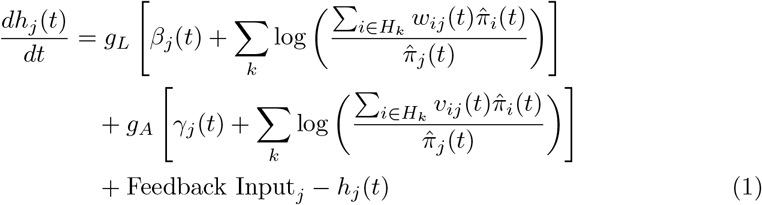

Here, Feedback Input_*j*_ refers to the summed input that unit *j* receives from *other* modules. Our implementation of such *inter-modular connections* is outlined further below.

### Activity Normalization

The output activity 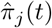 of each unit is computed via a softmax function, modelling lateral inhibition within hypercolumns:

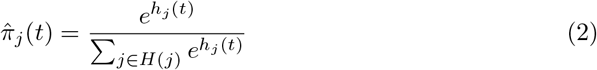

### Hebbian Learning

Hebbian learning updates excitatory synaptic weights *w*_*ij*_(*t*) and biases *β*_*i*_(*t*) based on running averages of unit activities. A ‘minimal background activity’ parameter *λ*_0_ ensures numerical stability and prevents extreme weight values in the absence of input.

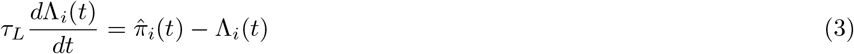

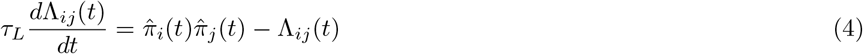

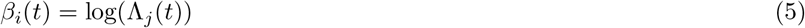

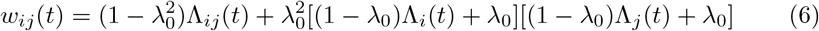

### Adaptation Mechanism

An analogous process governs inhibitory weights *v*_*ij*_(*t*) and biases *γ*_*i*_(*t*), which are updated on a slower timescale (governed by *τ*_*A*_):

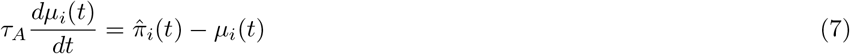

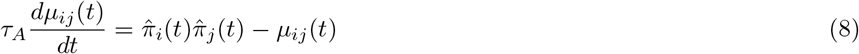

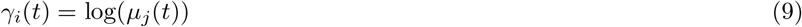

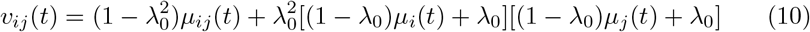

### kWTA Network

A k-Winner-Takes-All (kWTA) network is a variation on the BCPNN architecture described above. Unlike the BCPNN, the kWTA network does not organize its units into hypercolumns. Instead, the entire network competes globally, with only the *k* most active units at each time step producing non-zero activity. This approach, adopted from Fiebig & Lansner [16], more strictly enforces sparse network activity using a ‘hard WTA’ constraint.

On each time step, every unit computes its net excitatory input *h*, which evolves as follows:

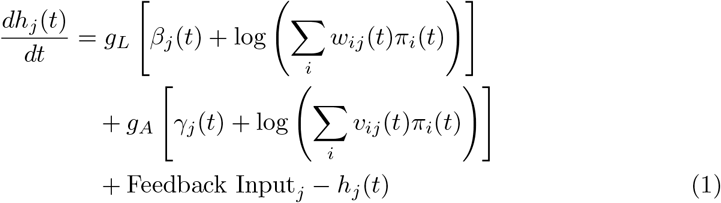

Here, *w*_*ij*_(*t*) and *β*_*j*_(*t*) represent the excitatory synaptic weights and biases, while *v*_*ij*_(*t*) and *γ*_*j*_(*t*) represent the inhibitory weights and biases. As previously, the terms *g*_*L*_ and *g*_*A*_ are scaling factors controlling the relative contributions of excitatory and inhibitory inputs.

### Activity Update: Hard WTA Rule

Unit activities in a kWTA network are binary (0 or 1) and determined using a hard WTA rule. At each time step, the *k* units with the highest *h*_*j*_(*t*) values are assigned an activity of 1, while the rest are set to 0:

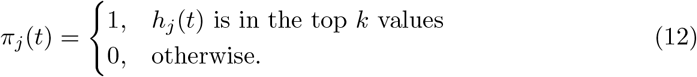

Here, *k* is given as the product of the total number of units (*N*) and a sparsity parameter (*α*). The kWTA network implements Hebbian learning and an adaptation mechanism using the same updating equations (3) to (10) as described for the BCPNN.

### Binary Module

Both BCPNNs and kWTA Networks effectively serve the purpose of auto-associatively storing and remembering their own activity patterns. This key feature makes them suitable for the CTX, HIP and BA_N_ modules of our model. In contrast, our model’s BA_P_ and BA_I_ modules can be conceptualized as populations of – non-interacting, for simplicity – cells that can be *recruited* by certain activity patterns of *other* modules. If a context is assumed to be encoded by a fixed activity pattern in BA_N_, then *associating valence* with that context is equivalent to *strengthening plastic synapses* from that BA_N_ ensemble to a selection of units in BA_P_ and/or BA_I_. BA_P_ and BA_I_ are implemented as ‘Binary Modules’.

The Binary Module operates by comparing the net input *h*_*j*_(*t*) received by each unit *j* to a fixed firing threshold *θ*. Units are activated (assigned a binary activity of 1) if their net input exceeds this threshold. The dynamics of net input and activity update are given by:

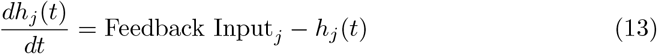

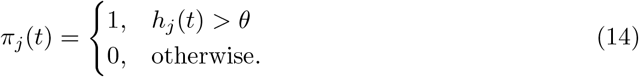

Hence, units of these modules receive their input entirely from other modules. This architecture enables BA_P_ and BA_I_ to encode valence-related quantities by recruiting varying numbers of active units.

### Inter-Modular Connections

In the sections above, we wrote Feedback Input_*j*_ to refer to the net input received by unit *j* from modules other than the one to which that unit belongs. The transmission of inputs from a *source* to a *target* module works very similarly to the transmission of excitatory inputs within a BCPNN or kWTA module.

### Updating Feedback Inputs

The feedback input to a target unit *j* is computed as a weighted sum of the activity of units in the source module, normalized and passed through a logarithmic activation. For each target unit *j*, the feedback input is given by:

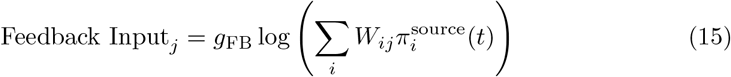

Here, *W*_*ij*_ represents the synaptic weight between source unit *i* and target unit *j*, and 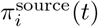 is the output activity of source unit *i* at time *t*. The term *g*_FB_ is a gain parameter controlling the contribution of the feedback input.

If inter-modular connections from multiple *source* modules are defined for a single *target* module, the contributions of these connections are summed.

### Hebbian Learning for Inter-modular Weights

Inter-modular connection weights *W*_*ij*_ adapt according to a Hebbian learning rule, driven by joint activity estimates between the source and target units:

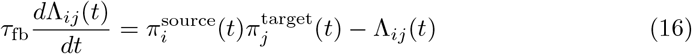

Here, Λ_*ij*_(*t*) is a running average that estimates the joint activity of source unit *i* and target unit *j*. These estimates are updated continuously over time with a timescale governed by *τ*_fb_.

The weights *W*_*ij*_(*t*) are computed from these activity estimates:

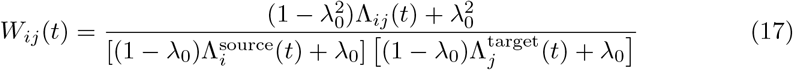

Here, 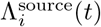 and 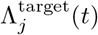 are marginal activity estimates of the source and target units, respectively – which are computed by the source and target *modules*, as per equation (3). As previously, the background activity parameter *λ*_0_ ensures numerical stability and prevents weights from diverging when activity levels are very low.

### Hyperparameters

Having defined the technical building blocks of our model, we provide the concrete parameter values used in the simulations. Synaptic gains (*g*) and learning time constants (*τ*) vary across the model’s *Perception, Sleep*, and *Recall* modes, as discussed in the main text.

**Table S1.** List of all modules in the model, their type, and number of units. The Sensory Cortex (SC) is special in that its activity is always manually set to a pattern encoding the current environment – or silenced during *Sleep*. Hence it does not actually behave like a BCPNN and effectively has no internal weights.

| Module Name | Module Type | No. Units | Comment |
| --- | --- | --- | --- |
| $SC$ | BCPNN | 500 | Activity manually set. |
| $EC_{IN}$ | Binary | 500 | cf. Fig S1. |
| HIP | kWTA | 350 |  |
| $EC_{OUT}$ | Binary | 500 | cf. Fig S1. |
| CTX | BCPNN | 500 |  |
| $BA_N$ | kWTA | 500 | |
| $BA_P$ | Binary | 250 | |
| $BA_I$ | Binary | 250 | |

**Table S2.** List of all plastic inter-modular connections in the model, as per Figure 1.

| Source Module | Target Module |
| --- | --- |
| HIP | $EC_{OUT}$ |
| HIP | CTX |
| CTX | $BA_N$ |
| HIP | $BA_N$ |
| $BA_N$ | $BA_P$ |
| $BA_N$ | $BA_I$ |

During *Recall*, it is always the case that *either* the gain *g*_*F B*_ of the *HIP → BA*_*N*_ connection *or* of the *CTX → BA*_*N*_ connection equals 1.0 – and the other equals 0.0. This depends on whether *HIP* successfully retrieved a memory (cf. Figure S1) – when this is not the case, ‘control’ over BA_N_ defaults to CTX.

**Table S3.**
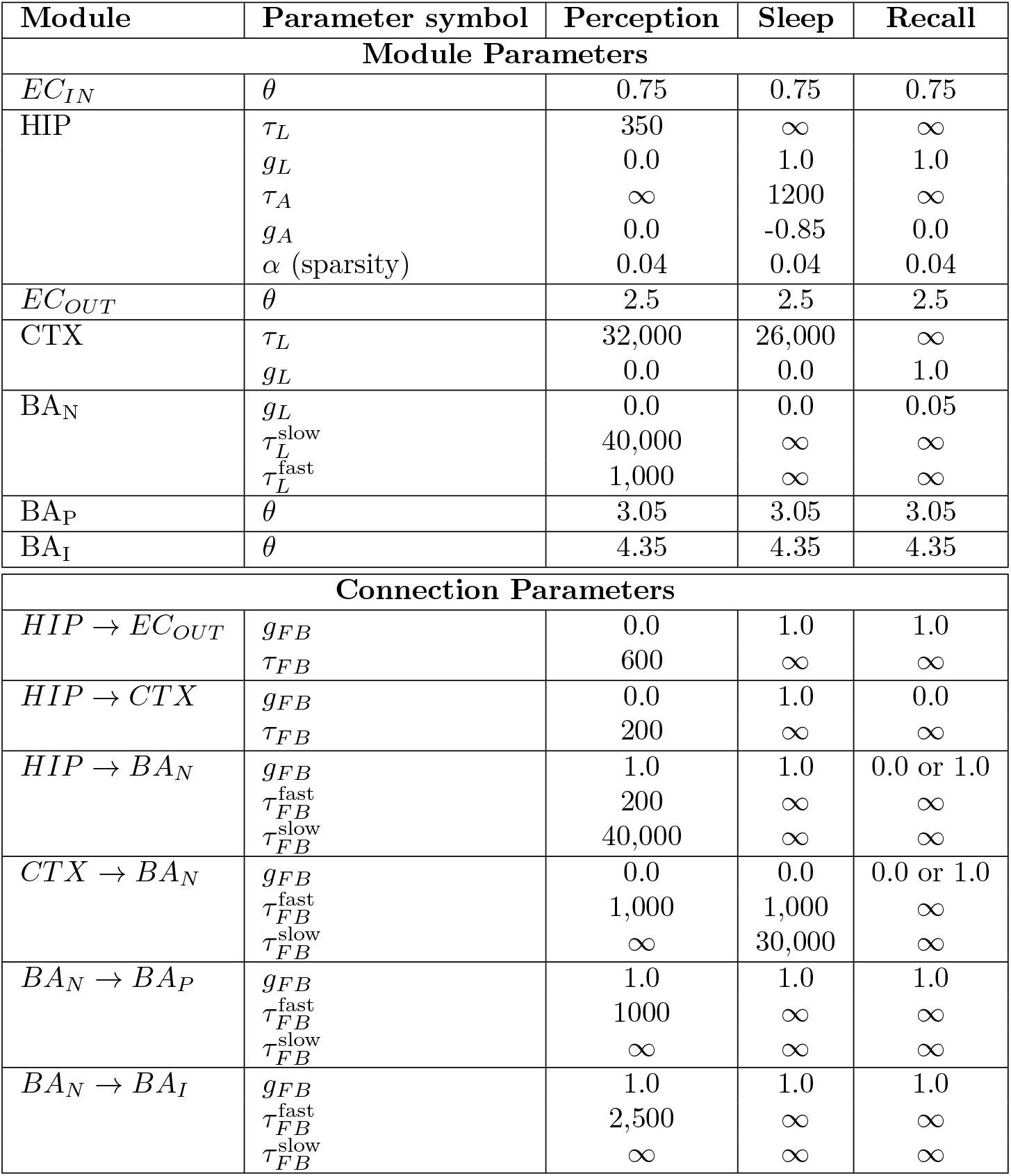
Module and connection parameters during Perception, Sleep, and Recall phases. Module parameters are listed above the double horizontal line, and inter-modular connection parameters are listed below. Parameter *θ* refers to a binary module’s firing threshold. *τ*_*L*_, *τ*_*A*_, and *τ*_*F B*_ denote the learning time constants for recurrent excitatory, inhibitory, and inter-modular weights, respectively. *g*_*L*_, *g*_*A*_, and *g*_*F B*_ represent the conductance (gain) of the respective synapses.

## S2 Appendix Full Update Cycle of the Model

This section describes the step-by-step update process of the model during a single simulation cycle. It outlines the sequence in which modules and connections are updated, to provide a clearer picture of how the model operates. The underlying Python code is provided in the supplementary materials.

### Stage 1 – Updating the Context-encoding Modules(HIP, CTX, BA N)

If the model currently receives an environmental input (`sensory_input` – an abstract, binary vector of length 500) – the activity of the engram modules is updated based on this input and pre-defined connection weight matrices. These feedforward connections are constant over time and are initialized randomly for any instance of the model. 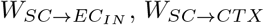 and *W*_*SC→BA N*_ are set to random permutations of the identity matrix. Entries of 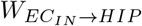 are drawn uniformly at random from [0, 1].

The following sequence occurs:

1. **Sensory Cortex Activity**: The output of the Sensory Cortex module (SC), *π*_*SC*_, is manually set to the provided <monospace>sensory_input</monospace>.
2. *EC*_*IN*_ **Activity**: Sensory Cortex activity is forwarded to *EC*_*IN*_. Each unit in *EC*_*IN*_ receives a binary input given by 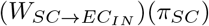 and attains it as its output to propagate the signal to HIP.
3. **Hippocampus (HIP) Activity**: The net input received by any unit in HIP is given by 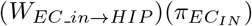. The sparsity No. units = *α N* = 0.04 350 = 14 most strongly innervated units become active, as per equation (12). Note that previously strengthened, recurrent synapses in HIP play no role at this point, as HIP activity during *memory formation* should be driven by external inputs rather than existing memories [129].
4. *EC*_*OUT*_ **Activity**: If the model is in **Perception** mode, the activity pattern of *EC*_*IN*_ is copied over to *EC*_*OUT*_ (cf. Figure S1).
5. **Cortex (CTX) and Amygdala (BA**_**N**_**) Activity** : Similar to step 3, the units of CTX and BA_N_ receive their input via (*W*_*SC→CT X*_)(*π*_*SC*_) or 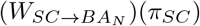 and attain their output as per equations (2) or (12), respectively.
6. **Within-Module Learning**: Now that the output of HIP, CTX and BA_N_ reflects the current environmental input, recurrent weights and related parameters of the modules are updated to support storing these activity patterns – unless the corresponding learning time constant *τ*_*L*_ is set to *∞*. These updates occur via equations (3) to (10) above.

If, on the other hand, no environmental input is present (say, during *Sleep*), the sequence of steps is shorter.

1. Update HIP, *EC*_*OUT*_ and CTX, in that order, as described by equations (1) to (12).
2. **Assessing validity of HIP Activity**: If the model is in **Recall** mode, an F1 Score is computed between the binary activity vectors of *EC*_*IN*_ and *EC*_*OUT*_. This measures whether *HIP* has *recalled* an engram that, during memory formation, was tied to an *EC*_*OUT*_ pattern that matches the current *EC*_*IN*_ activity. When this F1 Score exceeds 0.5, we say that HIP has successfully recalled a memory matching the current context. Then, gain *g*_*F B*_ of the *HIP → BA*_*N*_ connection is set to 1.0 and that of the *CTX → BA*_*N*_ connection is set to 0.0. Otherwise, the opposite occurs (cf. Fig S1).
3. Update BA_N_, as described by equations (1) to (10).

In summary, in stage 1 of the update cycle, environmental inputs are passed to the context-encoding modules HIP, CTX and BA_N_. Learning rate hyperparameters permitting, the resulting activity patterns are stored via Hebbian learning for future re-activation. When no environmental inputs are present (*Sleep* or *Recall*), the modules’ activities are determined by previously learned recurrent weights; new learning may yet occur during *Sleep*, as per Table S3. Note that, in *Perception* mode, the model always receives an environmental input.

### Stage 2 – Activating a Fear Response and Inter-modular Plasticity

Once a representation of the current – or currently recalled – context has been established in HIP, CTX and BA_N_, the model must activate a fear response and/or change the amount of fear associated with that context, as appropriate.

This occurs as follows:

1. The *recruitability* of all units in BA_P_ and BA_I_ is updated – according to a procedure which, for brevity, is described in a subsequent section of the appendix.
2. The activity of the model’s fear-evoking and -inhibiting cells, BA_P_ and BA_I_, is determined depending on plastic inputs arriving from BA_N_, as per equations (13) and (14).
3. CeA – the single unit encoding the model’s current fear response – receives a net input *h*_*CeA*_ equal to the number of active units in BA_P_, minus the number of active units in BA_I_. Its output, *π*_*CeA*_ is set to

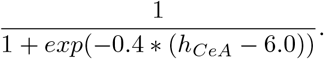

A sigmoid function is applied to *CeA*’s input to ensure that its output lies between 0 and 1 – the range of US strengths in our simulations.
4. The activity of the model’s *U-cell, π*_*U*_ is set to the magnitude of the US signal currently delivered by us, the experimenter. Whenever no US is delivered, *π*_*U*_ remains 0.
5. The activity of the model’s *A-cell*, intended to serve as a measure of *acute stress*, is updated as follows: In words, the stress level (*π*_*A*_) adjusts towards the US level (*π*_*U*_). Stress is assumed to rise more quickly when a US newly occurs or is strengthened than it falls when a US ceases or is weakened.

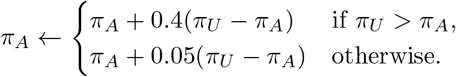
6. The difference Δ = *π*_*CeA*_ − *π*_*U*_ is recorded. We interpret it as a signed prediction error for the strength of the currently observed US.
7. If the absolute value of the prediction error, |Δ|, exceeds 0.3, the learning time constants *τ*_*L*_ (or *τ*_*F B*_) of the recurrent BA_N_ synapses and of the plastic *HIP → BA*_*N*_ and *CTX → BA*_*N*_ connections are set to their *fast* values 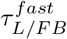 in Table S3). Else, the *slow* values are chosen.
8. If Δ *<* −0.05, i.e. if *π*_*U*_ *> π*_*CeA*_ + 0.05, the learning time constant of the *BA*_*N*_ *→ BA*_*P*_ connection is set to its *fast* value, 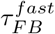, *divided* by |Δ|. This allows fear-evoking units in BA_P_ to be recruited – and more quickly so when a *very* unexpected US occurs. The learning time constant of *BA*_*N*_ *→ BA*_*I*_ is set to its *slow* value. Vice versa, if Δ *>* 0.05, the learning time constant of *BA*_*N*_ *→ BA*_*I*_ is set to 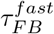 divided by Δ whereas that of *BA*_*N*_ *→ BA*_*P*_ is set to 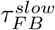 (cf. Table S3).
9. Now that the outputs of all of the model’s units have been set, learning takes place for all inter-modular, plastic synapses (cf. Table S2), as per equations (16) and (17).

An important particularity is necessary to allow the (‘Non-Hebbian’) recruitment of valence-coding units that are not yet active. When updating the *BA*_*N*_ *→ BA*_*P*_ and *BA*_*N*_ *→ BA*_*I*_ connections, equation (16) uses the activity 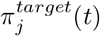 that the respective P- or I-cell *would have* if its net input were increased by the term

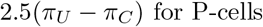

Or

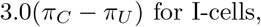

multiplied by that unit’s current *recruitability* value.

### Stage 3 – Sleep Homeostasis

The third and final part of the model’s update cycle consists of its sleep homeostasis mechanism – which we motivated and described in the main text of this paper.

The following steps are carried out:

1. If the *A-cell* activity, *π*_*A*_ lies above *T*_*stress*_ = 0.9, the *extinction threshold, A*_*P*_, of the homeostasis rule acting on BA_P_ is lowered from its default value 0.13 to 0.08 (favouring the long-term recruitment of fear-evoking units, as outlined below). The value of *T*_*stress*_ was chosen so that this putative stress effect only occurs after the delivery of extremely strong USs over several time consecutive time steps – such as in the experimental SEFL paradigm [130].
2. If the value of *A*_*P*_ currently lies below its default value of 0.13, it is incremented by 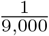. A full recovery of *A*_*P*_ thus occurs after (0.13 − 0.08) *\** 9, 000 = 450 time steps – roughly corresponding to two full day-night cycles in our simulations.

If the model is not currently in *Sleep* mode, the update cycle ends at this point. Otherwise, homeostatic changes are applied to synapses onto valence, coding amygdala cells, as follows:

1. Define *W*_*P*_ to be the weight matrix of the plastic *BA*_*N*_ *→ BA*_*P*_ connection, restricted to those rows that correspond to units in BA_N_ that are currently active.
2. Update the weights in *W*_*P*_ as per the cubic homeostasis rule

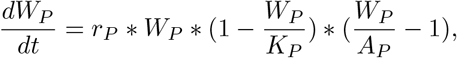

where *A*_*P*_ is as described above, and where *r*_*P*_ = 0.001, *K*_*P*_ = 0.31. Synaptic weights that would fall below 0 are set to 0 instead. The larger the gap between *A*_*P*_ and *K*_*P*_, the more stable the recruitment of P-cells by context-coding coding BA_N_ units.
3. To ensure that future updates to these synaptic weights will correctly account for these homeostatic changes (cf. Equation (16) above), the values of the ‘running average’ Λ_*ij*_ must be updated by inverting equation (17), as follows:

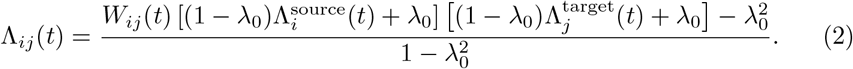

The meaning of the involved symbols is as described for equation (17).
4. Repeat the above steps 1 to 3, but for BA_I_ rather than BA_P_, with *r*_*I*_ = 0.001, *A*_*I*_ = 0.6125, *K*_*I*_ = 0.8.

### Recruitability of P- and I-cells

As presaged in Stage 2, Step 1 of the above update cycle, here we provide a formal definition of the way that the ‘recruitability’ of valence-coding amygdala (BA_P_ and (BA_I_) cells evolves in our model. As per Stage 2, Step 9, the current recruitability of each cell influences the likelihood of it being recruited into a context representation in case of a conditioning event. Computationally, the point of including such a parameter was to limit the amount of fear/extinction generalisation in the model – if conditioning in different environments always involved the same valence-coding cells, plasticity on the relevant synapses would quickly saturate.

The concept of a temporally evolving, cell-intrinsic *bias* affecting the allocation of neurons to newly formed activity patterns has previously been studied in the biological brain [131, 132], as well as in silico [52]. Yet, there is no clear consensus on how such a bias is likely to ‘behave’. We thus had a lot of freedom in our implementation, but formulated a list of design targets to guide us:

1. Each cell alternates between periods of high and low recruitability.
2. At any given time, only a relatively small fraction of valence-coding cells should be ‘highly recruitable’, to prevent a saturation of synapses (cf. above).
3. Groups of valence-coding cells that are simultaneously highly recruitable are broken up over time, so that – eventually – any combination of valence-coding cells may potentially appear in an engram. This is highly putative but, e.g., counteracts fear sensitization that would occur if the exact same group of P-cells was recruited in two contexts.
4. High-recruitability periods last approximately 200 time steps. This time window is longer than most conditioning sessions, but shorter than a day-night cycle, in our simulations. Valence-coding cell ensembles recruited into different context engrams on the same day are thus likely to overlap, such as to *link* the resulting associative memories [52].
5. The total recruitability of the system remains relatively stable over time.
6. Recruitability is a continuous quantity, with the full range of possible values being present in the population at any point in time. This, e.g., allows the recruitment of *few* cells in case of significant-but-weak predictions.

Formally, the recruitability of the fear-evoking P-cells, of which there are 250, evolves as follows:

1. When the model is first initialized, a small subset of P-cells is selected, independently at random, using a binomial distribution with *p* = 0.05. Each selected P-cell receives a ‘starting phase’ *θ*, again independently at random, uniformly from the interval [0, *π*]. For each remaining P-cell, *θ* is drawn uniformly from [−*π*, 0]. Then, each P-cell receives its initial recruitability, computed as

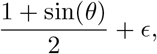

where the *ϵ*-term denotes random noise, drawn from 𝒩 (*µ* = 0, *σ* = 0.2), truncated from the right at *ϵ* = 0.8. Negative recruitability values are set to 0.
2. On any subsequent time step, the *phase* of each P-cell whose *current* recruitability exceeds 0.5 is advanced by 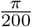. The duration for which cells remain *highly* recruitable is thus tightly regulated. On the other hand, the phase of each *remaining* P-cell has a (1*/*16)-chance of being advanced by 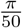. The duration for which cells are *unrecruitable* or ‘passive’ thus varies randomly – serving design target 3.) above. Recruitability values are again computed as 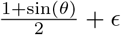.

The recruitability of the fear-inhibiting I-cells behaves independently from P-cells, but in exactly the same way. See Supp. Fig S2 for a visualization of the rule here discussed.

## S3 Appendix Simulation Protocols

**Table S4.**
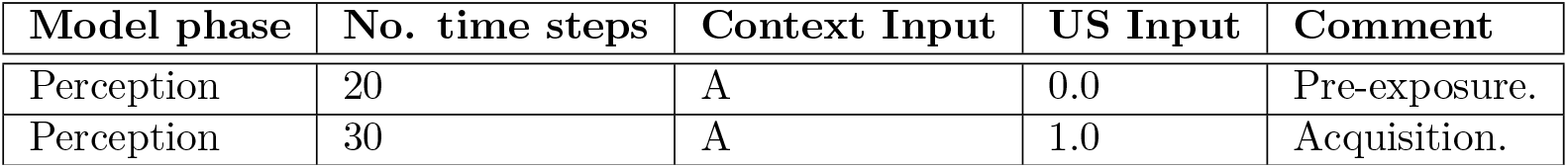
Demonstrating engram formation. Fig 4a.

| Model phase | No. time steps | Context Input | US Input | Comment |
| --- | --- | --- | --- | --- |
| Perception | 20 | A | 0.0 | Pre-exposure. |
| Perception | 30 | A | 1.0 | Acquisition. |

**Table S5.** Demonstrating sleep replay. Fig 4b.

| Model phase | No. time steps | Context Input | US Input | Comment |
| --- | --- | --- | --- | --- |
| Perception | 30 per context | 1 to 5 | 0.0 | – |
| Perception | 30 per context | 6 to 10 | 0.5 | Mild US to engage BA |
| Sleep | 165 | – | 0.0 | – |

**Table S6.** Assessing recall performance. Fig 4c. To simulate a scenario where some contexts are more likely to be remembered than others, the presentation lengths of the three contexts shown on any day are drawn from a Dirichlet-Multinomial distribution [133] (*n* = 60, *k* = 3, *α* = 20). To engage the BA_N_ module, every input is paired with a moderate US whose strength is drawn uniformly from 0.3 to 0.7. During *Recall* of a context, the original input pattern is presented for a single time step, with no further input over the 29 steps that follow. For each of HIP, CTX and BA_N_, a recall distance *d* is computed by comparing the module’s current output vector **u** to the activity **v** it had when the context was first presented during *Perception*. Recall is successful if 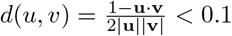[64].

| Model phase | No. time steps | Context Input | US Input | Comment |
| --- | --- | --- | --- | --- |
|  |  |  |  | Repeated 25 times |
| Perception | Cf. caption. | Three new contexts. | Cf. caption. | – |
| Sleep | 165 | – | 0.0 | – |
|  |  |  |  | Ultimately: |
| Recall | 30 per context. | Cf. caption. | 0.0 | – |

**Table S7.** Basic ‘AA’ fear extinction. Fig 4d.

| Model phase | No. time steps | Context Input | US Input | Comment |
| --- | --- | --- | --- | --- |
| Perception | 40 | A | 1.0 | Acquisition. |
| Perception | 100 | A | 0 | Extinction. |

**Table S8.** Effect of homeostasis on P-cell recruitment strength. Fig 6.

| Model phase | No. time steps | Context Input | US Input | Comment |
| --- | --- | --- | --- | --- |
| Perception | 40 | A | 1.0 | Acquisition. |
| Perception | 40 | B | 0 | No conditioning. |
| Sleep | 165 | - | 0 | Allowing homeostasis. |
| Recall | 1 | A | 0 | - |
| Recall | 1 | B | 0 | - |

**Table S9.** ‘ABA’, ‘ABC’ and ‘AAB’ fear renewal protocols. Fig 5b. In each case, contexts ‘B’ and ‘C’ are generated to be quite similar to context ‘A’, fixing 70% of 1s in the respective input pattern to lie in the same positions and drawing the remaining 30% independently at random.

| Model phase | No. time steps | Context Input | US Input | Comment |
| --- | --- | --- | --- | --- |
|  |  |  |  | ABA Renewal: |
| Perception | 20 | A | 1.0 | Acquisition. |
| Perception | 200 | B | 0.0 | Extinction. |
| Perception | 30 | A | 0.0 | Renewal and re-extinct. |
|  |  |  |  | ABC Renewal: |
| Perception | 20 | A | 1.0 | Acquisition. |
| Perception | 200 | B | 0.0 | Extinction. |
| Perception | 30 | C | 0.0 | Renewal and re-extinct. |
|  |  |  |  | AAB Renewal: |
| Perception | 20 | A | 1.0 | Acquisition. |
| Perception | 200 | A | 0.0 | Extinction. |
| Perception | 30 | B | 0.0 | Renewal and re-extinct. |

**Table S10.** Increased generalisation as fear memory ages. On each of the 15 days over which the fear memory in question ages, three random contexts are presented, each accompanied by a US of random, generally moderate strength. Fig 5c.

| Model phase | No. time steps | Context Input | US Input | Comment |
| --- | --- | --- | --- | --- |
| Perception | 25 | A | 0.9 | Fearful context. |
| Perception | 25 | B | 0.0 | Similar context |
| Perception | 25 | C | 0.0 | Dissimilar conte |
| – | – | – | – | 15 ‘delay days’: |
| Perception | 30 | Random 1 | 0.8 * Beta(1.25,3) |  |
| Perception | 30 | Random 2 | 0.8 * Beta(1.25,3) |  |
| Perception | 30 | Random 3 | 0.8 * Beta(1.25,3) |  |
| Recall | 10 | A (on 1st step) | 0.0 |  |
| Recall | 10 | B (on 1st step) | 0.0 |  |
| Recall | 10 | C (on 1st step) | 0.0 |  |
| Sleep | 165 | – | – |  |

**Table S11.**
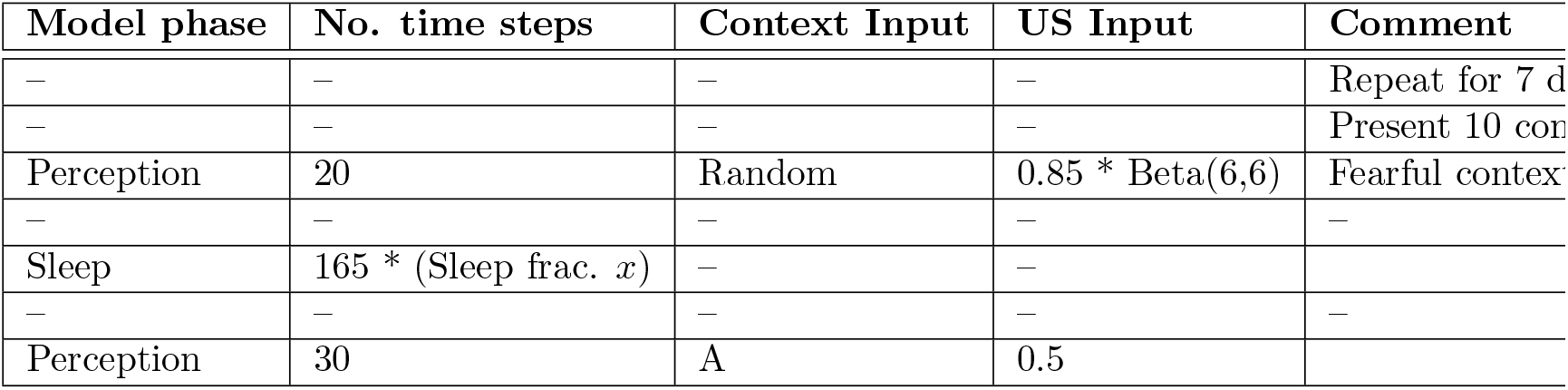
Sleep deprivation. We defined multiple instances of the model, each with a different sleep fraction *x*. On each of 7 simulated days, the models were presented 10 random contexts, accompanied by moderate US delivery, causing the formation of fear memories in some of them. Then, light fear acquisition was performed in context A. At the end of training, the fear response acquired by each model was recorded. Fig 7.

| Model phase | No. time steps | Context Input | US Input | Comment |
| --- | --- | --- | --- | --- |
| – | – | – | – | Repeat for 7 d |
| – | – | – | – | Present 10 contexts |
| Perception | 20 | Random | 0.85 * Beta(6,6) | Fearful contexts |
| – | – | – | – | – |
| Sleep | 165 * (Sleep frac. $x$) | – | – | – |
| – | – | – | – | – |
| Perception | 30 | A | 0.5 | – |

**Table S12.** SEFL protocol. Exposure to harmless contexts ‘C’ and ‘D’ is included to ‘balance out’ hippocampal replay during the following *Sleep* phase. Fig **??**a and **??**d.

| Model phase | No. time steps | Context Input | US Input | Comment |
| --- | --- | --- | --- | --- |
| Perception | 30 | A | 1.0 (0.0 in controls) | Traumatic stress |
| Perception | 30 | C | 0.0 | Harmless. |
| Perception | 30 | D | 0.0 | Harmless. |
| Sleep | 165 | – | – | – |
| Perception | 3 | B | 0.9 | Moderate US. |
| Perception | 27 | B | 0.0 | – |
| Perception | 30 | C | 0.0 | Harmless. |
| Perception | 30 | D | 0.0 | Harmless. |
| Sleep | 165 | – | – | – |
| Recall | 30 | B (on 1st step) | 0 | - |

**Table S13.** SEFL protocol – Order reversed. Same as Table S12, but with the order of the ‘trauma’ and ‘moderate US’ components of the SEFL protocol reversed. Fig **??**b.

| Model phase | No. time steps | Context Input | US Input | Comment |
| --- | --- | --- | --- | --- |
| Perception | 3 | B | 0.9 | Moderate US. |
| Perception | 27 | B | 0.0 | – |
| Perception | 30 | C | 0.0 | Harmless. |
| Perception | 30 | D | 0.0 | Harmless. |
| Sleep | 165 | – | – | – |
| Perception | 30 | A | 1.0 (0.0 in controls) | Traumatic stress |
| Perception | 30 | C | 0.0 | Harmless. |
| Perception | 30 | D | 0.0 | Harmless. |
| Sleep | 165 | – | – | – |
| Recall | 30 | B (on 1st step) | 0 | - |

## S4 Appendix

### Supplementary Figures

**Table S14.** SEFL protocol – Delayed Sensitization test. Same as Table S12, but with 15 ‘delay days’, on each of which three random contexts (of moderate fearfulness) are presented. After those 15 days, ‘moderate fear acquisition’ in context ‘B’ is performed as in the default SEFL protocol. Fig **??**c.

| Model phase | No. time steps | Context Input | US Input | Comment |
| --- | --- | --- | --- | --- |
| Perception | 30 | A | 1.0 (0.0 in controls) | Traumatic s |
| Perception | 30 | C | 0.0 | Harmless. |
| Perception | 30 | D | 0.0 | Harmless. |
| Sleep | 165 | – | – |  |
| – | – | – | – | 15 ‘delay days’ |
| Perception | 30 | Random 1 | drawn from Beta(3,4) |  |
| Perception | 30 | Random 2 | drawn from Beta(3,4) |  |
| Perception | 30 | Random 3 | drawn from Beta(3,4) |  |
| Sleep | 165 | – | – |  |
| Perception | 3 | B | 0.9 | Moderate US |
| Perception | 27 | B | 0.0 | – |
| Perception | 30 | C | 0.0 | Harmless. |
| Perception | 30 | D | 0.0 | Harmless. |
| Sleep | 165 | – | – |  |
| Recall | 30 | B (on 1st step) | 0 | - |

**Fig S1.**
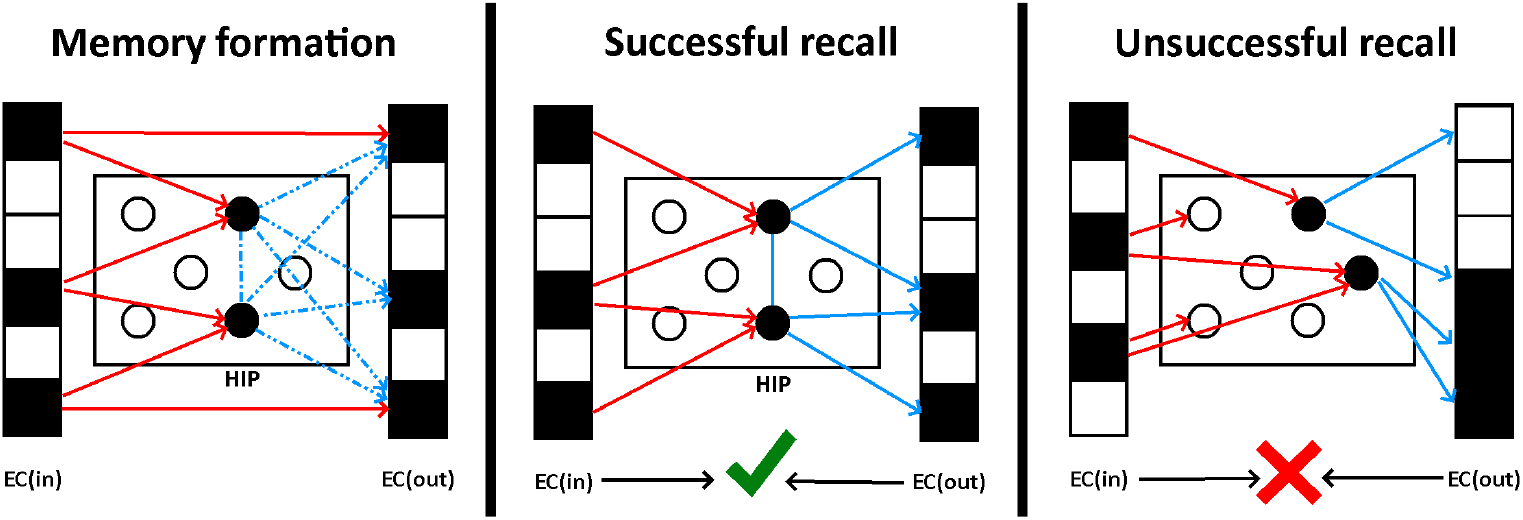
Memory formation and recall in HIP. Any engram formed in the HIP module becomes associated with a pattern in the EC output layer, equal to the input pattern that led to its formation. Dotted arrows denote plastic synapses being strengthened. When the engram is retrieved by the same (or a sufficiently similar) cue, the original input pattern is activated in the output layer and observed to match the current input.

**Fig S2.**
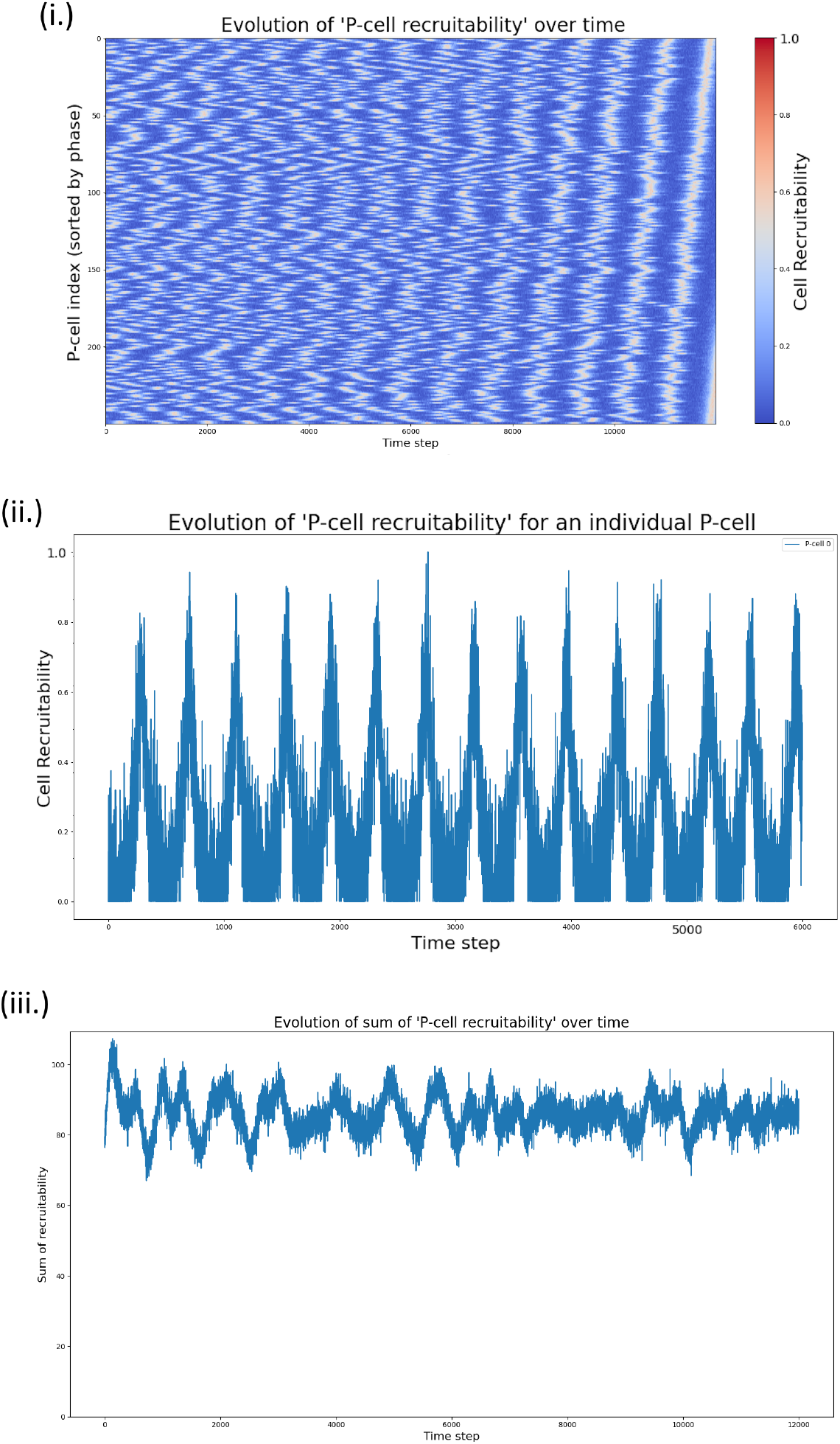
P-cell recruitability. **(i.)** Rows of the heatmap denote individual P-cells, columns denote time steps of our simulation. Rows of the heatmap were sorted according to the cells’ ‘phase’ (cf. below) after 12,000 time steps. The blurring of the white vertical lines at prior time points indicates that P-cells change the ‘partners’ they are highly recruitable *with* over time. **(ii.)** Evolution of the recruitability for an individual P-cell, corresponding to the first row of plot (i). Time windows of high recruitability last about 200 time steps and are interspersed by considerably longer phases of low recruitability. **(iii.)** The total sum of the recruitability values of all P-cells remains fairly stable over time.

In more detail: During *Perception*, the context-defining activity pattern of the EC input module (*EC*_*in*_) is *copied* to *EC*_*out*_, and synapses between HIP and *EC*_*out*_ are rapidly strengthened. During *Recall*, the direct projection from input-to output layer is switched off, but the gain of the HIP-to-*EC*_*out*_ connection is switched on. If a memory was recalled, *EC*_*out*_ should therefore contain the pattern that was present when that memory was formed, while the activity of *EC*_*in*_ depends on the current sensory input. An F1 score is computed between *EC*_*in*_ and *EC*_*out*_ pattern; if it exceeds a certain threshold, recalled and current context are said to match. If a mismatch is detected, inputs from HIP to BA_N_ are switched off and the responsibility of retrieving a BA_N_ representation for the current environment falls to CTX.

